# Mapping the functional importance of site-specific ubiquitination across the human proteome

**DOI:** 10.1101/2025.10.08.681129

**Authors:** Julian van Gerwen, Maximilian Fottner, Shengbo Wang, Bede Busby, Ellen Boswell, Paul Schnacke, Andrea C. Carrano, Malina A. Bakowski, Emily R. Troemel, Romain Studer, Marta Strumillo, Maria-Jesus Martin, J. Wade Harper, Kathrin Lang, Andrew R. Jones, Eric J. Bennett, Juan Antonio Vizcaíno, Inigo Barrio-Hernandez, Pedro Beltrao

## Abstract

Protein ubiquitination regulates cell biology through diverse avenues, from quality control-linked protein degradation to signaling functions such as modulating protein-protein interactions and enzyme activation. To date, hundreds of thousands of ubiquitination sites (ubi-sites) have been identified, however fewer than 1% have known functional roles. Here, we assembled a human reference ubiquitinome of 108,341 ubi-sites by harmonizing public proteomics data. To pinpoint critical regulatory events requiring ubiquitination at a precise site, we mapped ubi-site conservation across proteomics data from six non-human species. Perturbation proteomics revealed that highly conserved ubi-sites are more likely to regulate signaling functions rather than proteasomal degradation. To further prioritize site-specific ubiquitination relevant for organismal fitness, we constructed a machine learning-based positional importance score for more than 100,000 ubi-sites, which identifies sites regulating diverse protein functions and rationalizes genetic vulnerabilities. Finally, we employed chemical genomics to validate the functional relevance of high-scoring ubi-sites and leveraged genetic code expansion to demonstrate that ubiquitination of K320 in the RNA-regulator ELAVL1 disrupts RNA binding. Our work reveals systems-level principles of the ubiquitinome and provides a powerful resource for studying site-specific protein ubiquitination.

## Introduction

Ubiquitination is a reversible post-translational modification (PTM) critical for cell biology. It involves the attachment of the 76-amino acid protein ubiquitin to specific lysine residues either as a single moiety or polymeric chains, catalysed by the concerted activity of E1, E2, and E3 enzymes, and opposed by deubiquitinating enzymes^1–3^. The best appreciated consequence of ubiquitination is the targeting of aberrant proteins for proteasomal degradation by K48-linked polyubiquitin, but ubiquitination can also modulate protein-protein interactions, induce protein conformational changes, and alter protein subcellular location, among other functions^1,4^. These functions place ubiquitin as a central player both in protein quality control and diverse signaling processes including DNA repair^5^, cell cycle progression^6^, and endocytosis^7^, underscoring its importance for cellular homeostasis.

Many proteins contain multiple sites of ubiquitin attachment which may often be functionally redundant. Nevertheless, there is accumulating evidence of site-specific regulatory ubiquitination, where the mono or poly-ubiquitination of a specific lysine alters protein function to maintain cellular homeostasis. For instance, mono-ubiquitination of K523 and K561 in the FANCI-FANCD2 complex causes conformational changes that are critical for interstrand crosslink repair^8^, and mono and poly-ubiquitination of separate lysines on histone H2A recruit different effectors that drive distinct outcomes for DNA repair and transcriptional regulation^9^, among other examples^10–15^. Furthermore, while there is likely often redundancy between the ubi-sites that target a protein for proteasomal degradation by K48-polyubiquitination, there are examples of proteins where only one or a few lysines are primarily responsible for or capable of promoting degradation^16–23^. The distribution of such functionally important, position-specific ubiquitination events across the human proteome remains unknown.

Our understanding of the proteome-wide repertoire of ubiquitination sites has been transformed over the last two decades by mass spectrometry-based proteomics, which allows the unbiased identification and quantification of ubiquitination events, albeit without information on ubiquitin chain composition^24–26^. Pioneering proteomics studies quantified thousands of ubi-sites in response to diverse cellular perturbations, and identified that distinct sites within the same protein often display opposing dynamic profiles, alluding to site-specific regulatory behaviour^27–29^. However, the discovery afforded by proteomics has rapidly outpaced functional characterisation - of the 100,000 human ubi-sites now identified, only approximately 1,000 have an experimentally determined regulatory role, as reported in the PhosphositePlus repository^30^.

Here, we have addressed the system-wide dearth of functional characterisation for human ubi-sites with a focus on site-specific ubiquitination. We first assembled a high-confidence reference ubiquitinome of more than 100,000 ubi-sites by re-analyzing 11 published proteomics datasets. Leveraging published and novel ubiquitin proteomics data across six species and 103 cellular perturbations, we identified thousands of site-specific ubiquitination events conserved across genomes and protein domain families, and found that these sites tend to mediate non-degradative signaling functions rather than proteasomal degradation. We then constructed a positional importance score for 100,000 ubi-sites by integrating evolutionary, proteomic, and structural features using machine learning, in order to prioritise site-specific ubiquitination important for cellular homeostasis. High-scoring ubi-sites regulate protein function through diverse avenues and are frequently disrupted by pathogenic mutations. Finally, we used chemical genetics to identify cellular roles for orthologs of high-scoring ubi-sites in yeast, and employed genetic code expansion to demonstrate that ubiquitination of K320 in the human post-transcriptional regulator ELAVL1 inhibits RNA binding. Our work constitutes a significant advance in our understanding of the human ubiquitinome and a powerful resource for the study of ubiquitin site function.

## Results

### Constructing a human reference ubiquitinome

Mass spectrometry-based proteomics experiments routinely identify tens of thousands of ubi-sites^25,26,31^, which are collated in PTM-centric resources such as PhosphositePlus^30^. However, this approach can lead to inaccurate PTM identifications, if false discovery rates are not statistically controlled across multiple studies and datasets^32^. Hence, we first sought to generate a high-confidence reference set of human ubi-sites by systematically re-analyzing public mass spectrometry-based proteomics datasets (Fig. 1a).

**Figure 1:**
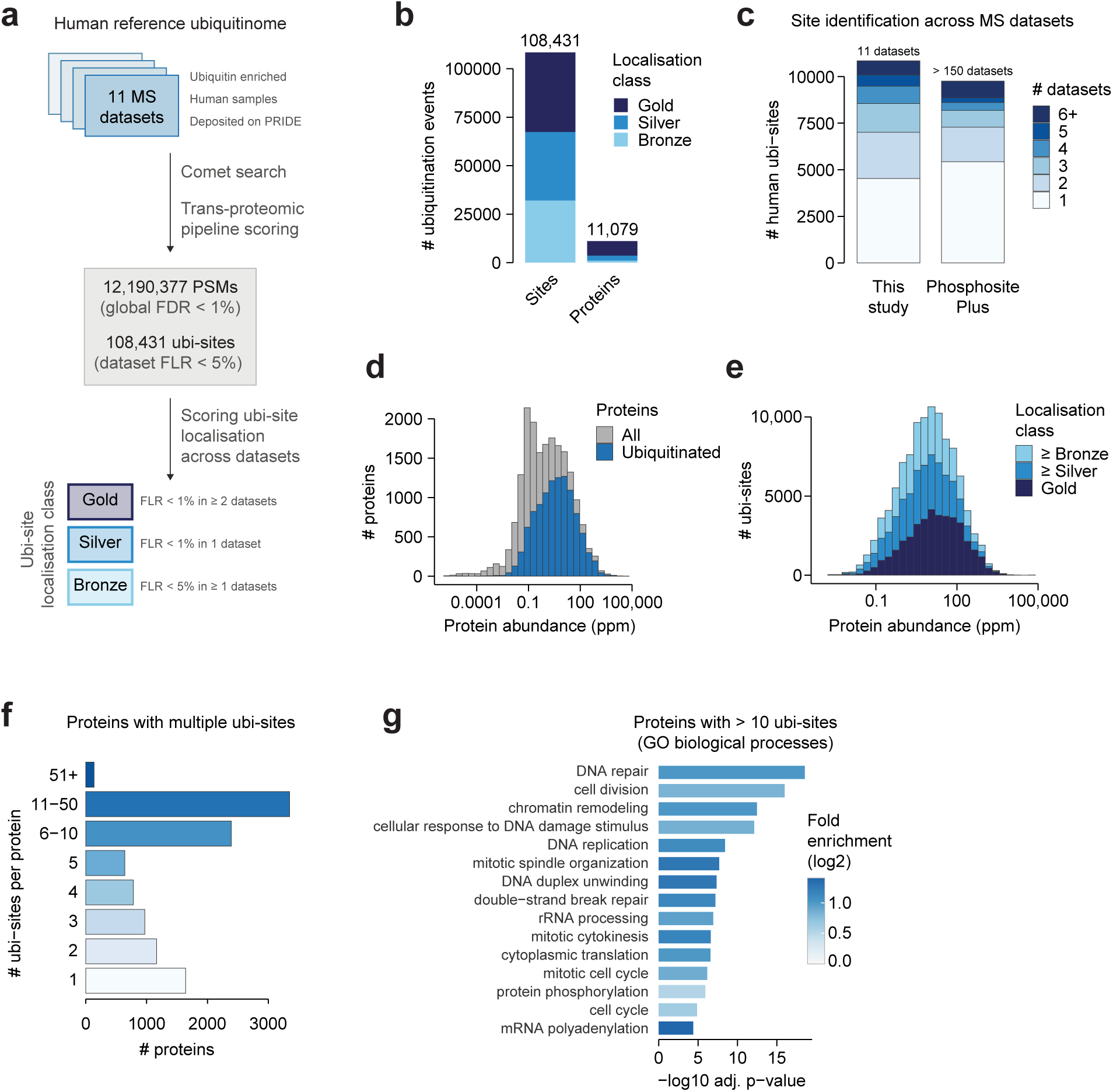
Constructing a human reference ubiquitinome A) Pipeline for the re-analysis of ubiquitin-proteomics spectral data and their aggregation into a reference ubiquitinome. B) The number of ubiquitinated sites and proteins in the human reference ubiquitinome, stratified by their confidence level. A protein’s confidence level is calculated as the maximum across all of its sites. C) The number of human ubiquitination sites in our reference ubiquitinome and PhosphositePlus^30^, stratified by the number of datasets in which each site is quantified. D) The abundance of all proteins in the human reference ubiquitinome compared to the entire proteome, using abundance measurements from PaxDB^45^. E) The protein abundance of ubiquitination sites at different levels of identification confidence. F) The number of identified ubi-sites per protein. G) The top 15 significantly overrepresented GO biological process terms among proteins with more than 10 ubi-sites, relative to all proteins in the human reference ubiquitinome (one-sided Fisher’s exact test, Benjamini-Hochberg p-value adjustment).

We began by curating 11 recent ubiquitin proteomics datasets from the PRIDE data repository^33^ that covered a range of cell types and experimental conditions (Table S1^26,31,34–42^). We then extracted and re-analyzed raw proteomics data using both community-standard and bespoke analysis tools (see Methods), resulting in a total of 12,190,377 peptide-spectrum matches (PSMs) at 1% global false-discovery rate per dataset, 108,341 ubi-sites at 5% false-localisation rate per dataset, and 11,079 ubiquitinated proteins (Fig. 1a-b). Using our established scoring scheme for PTM identification confidence^43^ we found that 41,070 ubi-sites displayed multiple high-stringency identifications (FLR < 1% in ≥ 2 datasets, “Gold” confidence) and 35,469 ubi-sites displayed one high-stringency identification (FLR < 1% in 1 dataset, “Silver” confidence), underscoring the reliability of our pipeline (Fig. 1a-b).

Our ubi-site identifications also showed strong overall concordance with both PhosphositePlus (71,268 shared sites) and PTMAtlas^44^, a recent database generated from reanalysis of public proteomics data (75,930 common sites, Fig. S1a). Furthermore, our dataset likely suffers from fewer false positive identifications than PhosphositePlus, since PhosphositePlus contains a greater fraction of sites identified in only a single contributing dataset, despite aggregating many more datasets than us (more than 150 published and internal datasets). Finally, ubiquitinated proteins in our dataset spanned almost the entire range of abundances in the human proteome with only minor differences compared to non-ubiquitinated proteins and across confidence levels (Fig. 1d-e^45^), indicating only a small influence of the bias of mass spectrometry to high-abundance molecules.

Our reference ubiquitinome displayed a wide range in the number of ubi-sites detected per protein (Fig. 1f). This was not strongly associated with protein abundance, protein length, or lysine content, suggesting this observation is not driven by biases in detectability by mass spectrometry or the availability of modifiable lysines, and instead represents a biological signal (Fig. S1b-d). The 3,486 proteins with more than 10 ubi-sites were enriched in DNA repair and cell cycle pathways (Fig. 1g) and at diverse cellular locations including the centrosome and spindle (Fig. S1e), consistent with previous work identifying these as focal points for extensive ubiquitin regulation^5,26^. Overall, we have generated a large-scale, high-confidence atlas of human ubiquitin sites that can provide rich insight into the human ubiquitination landscape. To ensure that these data meet FAIR guidelines^46^, the outputs have been made available in PRIDE (PRIDE number PXD068989, Table S2).

### Characterizing the conservation of human ubi-sites

As a first step to a functional understanding of the human ubiquitinome, it would be valuable to know which ubi-sites represent positionally important regulatory events - cases where the mono or poly-ubiquitination of a specific lysine is required to tune protein function and balance cellular homeostasis. Conservation analysis is a powerful tool to identify such ubi-sites as it pinpoints protein modifications that have been positionally constrained by evolution to support organismal fitness^47–49^.

To assess ubi-site conservation, we assembled published and in-house ubiquitin proteomics data from six species spanning a range of evolutionary distances from *Homo sapiens* (Fig. 2a, Table S3 and S4^50–56^). After aligning protein sequences across species we classified human ubi-sites into five levels of conservation, where the lowest level represents lysines largely absent in other species, the highest levels represent lysines with ubiquitination detected in other species, and intermediate levels represent lysines that are conserved in other species but without measured ubiquitination, which still have the potential to be ubiquitinated (Fig. 2b). Human GAPDH demonstrates that one protein can display a range of ubi-site conservation (Fig. 2c).

**Figure 2:**
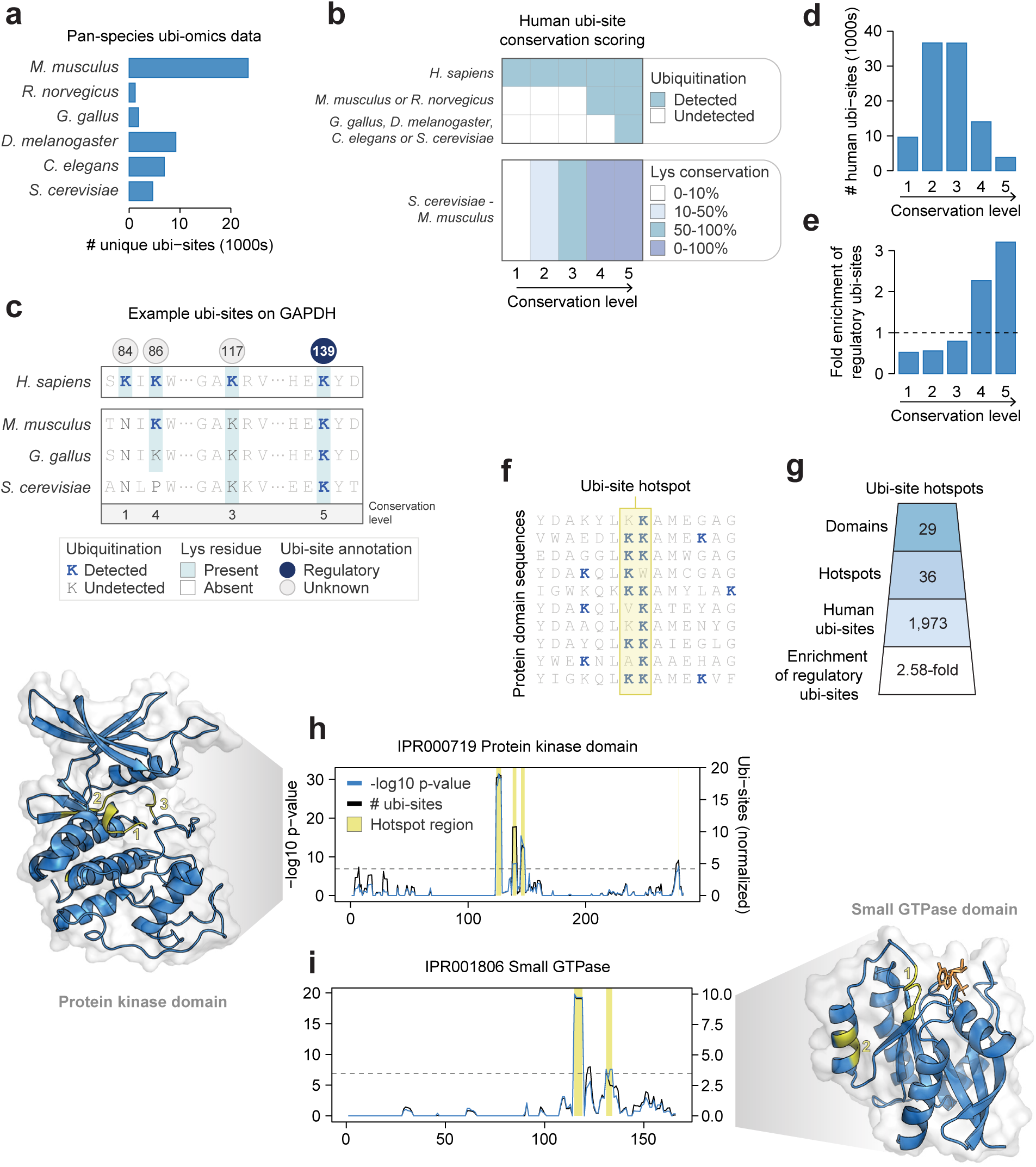
Characterizing the conservation of human ubi-sites A) The number of ubi-sites identified in a compilation of ubiquitin-proteomics data across six non-human species. B) Rules for scoring the conservation of human ubi-sites based on the detection of ubiquitin and the presence of lysine at the orthologous position in other species. Lys conservation indicates the percentage of residues at the given alignment position that are lysine. C) Conservation of several ubi-sites in GAPDH including regulatory ubi-site annotations from PhosphositePlus. D) The number of human ubi-sites at each conservation level. E) The enrichment of ubiquitin sites annotated to perform non-degradation regulatory functions at each conservation level (PhosphositePlus^30^). F) Conceptual illustration of ubi-site hotspots: regions within aligned protein domain sequences enriched in ubiquitination. G) The number of identified ubi-site hotspots, hotspot domains, and human hotspot sites. The enrichment of human hotspot sites annotated to perform non-degradation regulatory functions (see Methods) is shown. H-I) Identification of ubi-site hotspots in two protein domains. The black line indicates the average number of ubi-sites observed across the domain sequence alignment within a rolling window, normalised by subtracting the number of ubi-sites expected by chance. The blue line indicates the p-value associated with the enrichment of ubi-sites at each alignment position. The horizontal line indicates a Bonferroni-corrected p-value cut-off of 0.01 (uncorrected p-value < 1.29×10^-7^). Positions with a-log10 p-value above this cut-off and average number of phosphosites per window higher than 2 are classified as hotspot regions and highlighted with a yellow bar. Hotspot regions are mapped onto representative structures for each domain in yellow. Multiple hotspots are labeled in their order in the domain sequence.

Ubi-site conservation was generally limited; of the 100,609 analyzed sites, only 17,805 displayed ubiquitination in *M. musculus* or *R. norvegicus* (conservation level 4), and 3,814 displayed additional ubiquitination in the more divergent species *G. gallus, D. melanogaster, C. elegans,* or *S. cerevisiae* (conservation level 5, Fig. 2d). This may underestimate true conservation, as our non-human proteomics data contains far fewer individual datasets and ubi-site identifications compared to human. Regardless, the two highest conservation levels were enriched for ubi-sites annotated with regulatory, non-degradation functions in PhosphoSitePlus (Fig. 2e, 2.27- and 3.21-fold enrichment for levels 4 and 5), and also showed enrichment - albeit to a lesser degree - for degradation-related functions (Fig. S2a, 2.00- and 2.00-fold enrichment for levels 4 and 5). For instance, K139 can stabilise and activate GAPDH by K63-linked polyubiquitination^57^, is the only annotated regulatory ubi-site on GAPDH in PhosphositePlus, and was conserved from humans to *S. cerevisiae* (level 5, Fig. 2c). This confirms that evolutionary constraint identifies functionally relevant modifications with known regulatory roles.

We next employed an orthogonal approach to identify conserved ubi-sites. Following our previous work on protein phosphorylation^49^ and related work by others^58,59^, we searched for small regions in protein domains that are repeatedly ubiquitinated within and across genomes at a rate higher than expected by chance, which we term “ubi-site hotspots” (Fig. 2f, see Methods). Such modifications have likely been conserved to regulate domain functions for millions of years of protein evolution. We identified 36 ubiquitination hotspots within 29 InterPro protein domains, comprising a total of 1,973 human ubiquitin sites (Fig. 2g, Table S5, Data S1). These hotspots were enriched in ubiquitin sites annotated with non-degradation functions (2.58-fold, one-sided Fisher’s exact test p-value = 0.00062) and to a lesser extent degradation functions (1.41-fold, one-sided Fisher’s exact test p-value = 0.23), demonstrating that domain-level conservation identifies functionally important modifications.

To further illustrate the value of ubi-site hotspot analysis, we highlight representative protein domains as case studies. We identified three ubi-site hotspots proximal to the activation loop of the eukaryotic protein kinase domain, which were independently identified in previous work^60^ (IPR000719, Fig. 2h). The most prominent of these hotspots contains ubi-sites in 163 of the approximately 500 human protein kinases^61^, including seven ubi-sites experimentally demonstrated to regulate their respective kinase (Table S5), pointing to a general mechanism controlling kinase signaling. Second, we identified two hotspots in the small GTPase domain (IPR001806, Fig. 2i). The most prominent hotspot is adjacent to the GTP/GDP-binding sites and mono-ubiquitination of HRAS K117 in this hotspot has been shown to promote nucleotide exchange^12^, highlighting its regulatory potential. Other illustrative examples include hotspots at the tip of the C2H2-type zinc finger domain (IPR013087), at the GEF-GTPase interface of the Dbl homology domain (IPR000219), and proximal to the ubiquitin-binding cleft in the catalytic domain of USP deubiquitinases (IPR001394, Fig. S3a-c). Based on the above findings, we anticipate that highly conserved ubi-sites and hotspot ubi-sites represent a wealth of site-specific regulatory events important for cellular homeostasis.

### Characterizing the regulation of human ubi-sites

Beyond conservation, the dynamic regulation of ubi-sites across cellular perturbations provides an orthogonal view into their functional and regulatory characteristics. To enable analysis of ubi-site regulation, we analyzed quantitative human ubiquitin proteomics data from 14 publications and generated an additional 10 proteomics datasets in-house (Fig. 3a, Table S3^27–29,62–72^). The resulting compilation encompasses 103 control-matched perturbation conditions, 62,094 human ubi-sites, and 452,191 quantified ubi-site fold changes (Fig. 3b). These perturbations include diverse cellular stressors such as DNA damaging agents and ER stressors, as well as inhibitors of deubiquitinases (DUBs), neddylation, or the proteasome (Fig. 3c).

**Figure 3:**
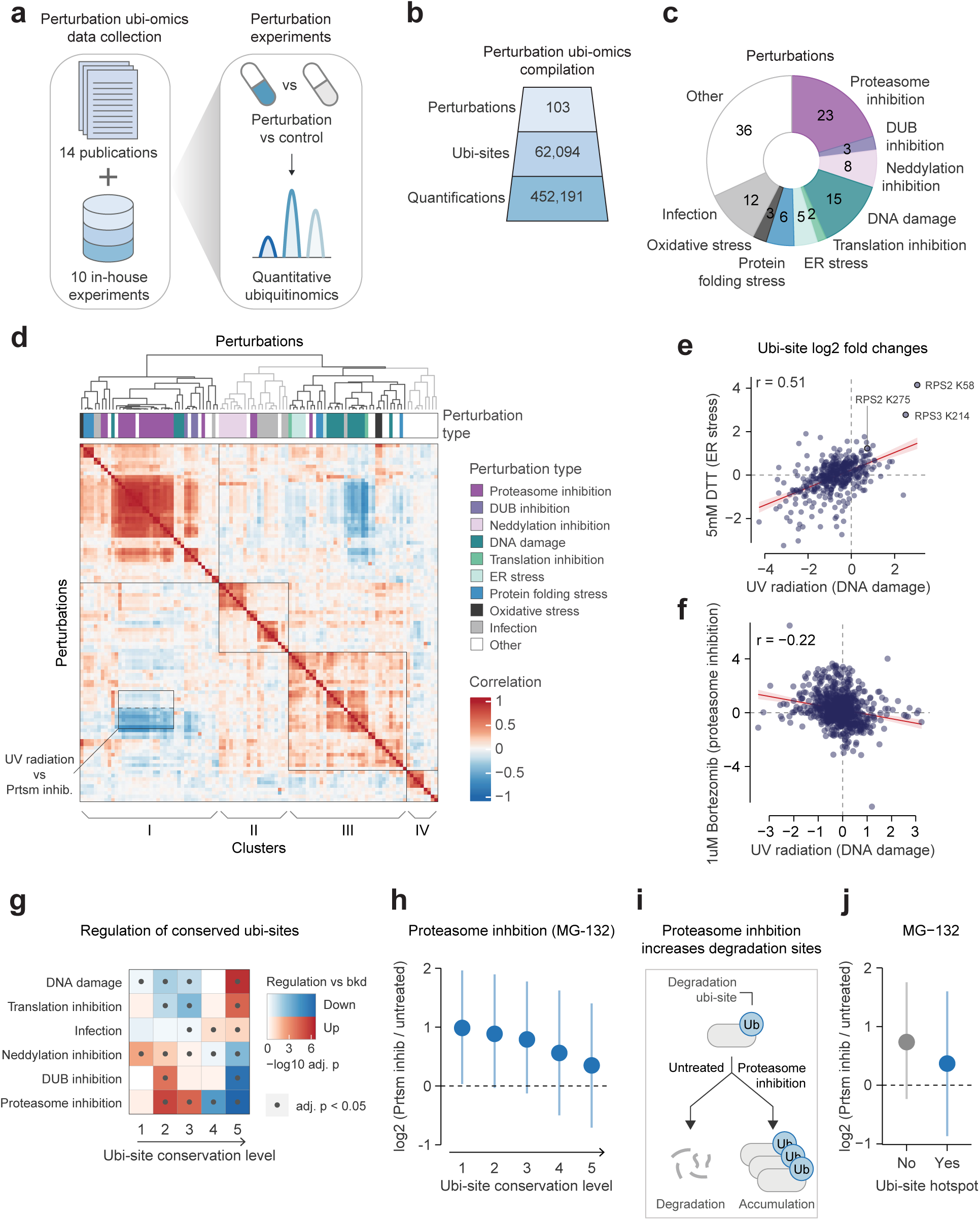
Characterizing the regulation of human ubi-sites A) Ubiquitin proteomics experiments measuring cellular perturbations in human cells were collated from 14 publications and 10 in-house experiments. B) The number of perturbation conditions, quantified ubi-sites, and quantified ubi-site fold change values across compiled ubiquitin proteomics data. C) The distribution of different types of perturbations in compiled ubiquitin proteomics data. D) Pairwise Pearson’s correlations between perturbation conditions based on ubi-site log2-fold changes. Perturbations were divided into four clusters by hierarchical clustering using Pearson’s correlation as the distance metric, indicated by different colours in the dendrogram and labels below the heatmap. The negative correlation between UV radiation conditions and proteasome inhibition conditions is outlined, where the bottom half of the box indicates UV radiation of cells with prior proteasome inhibition, while the top half indicates UV radiation of untreated cells. E-F) Examples illustrating E) a positive (UV radiation vs. ER stress) and F) negative (UV radiation vs. proteasome inhibition) correlations of ubi-site responses between perturbation types. Pearson correlation coefficients (r) are shown. Ribosomal ubi-sites implicated in the unfolded protein response are labeled^29^. G) Two-sided gene-set tests (Limma v3.56.2, geneSetTest, type = “t”, alternative = “either”) were performed to identify conditions in which ubi-sites at each conservation level were up-or down-regulated relative to all other ubi-sites. P-values were adjusted by the Benjamini-Hochberg procedure, converted into log10-values, and averaged within each perturbation group. Only perturbation groups with at least one significant enrichment (adj. p < 0.05) are shown. H) Mean log2-fold changes (± s.d.) of ubi-sites under proteasome inhibition (MG-132 treatment), grouped by evolutionary conservation level. Final values were obtained by normalising and averaging log2 fold-changes from multiple experiments (see Methods). The number of ubi-sites in each group from left-to-right are 1451, 6404, 7869, 5234, and 2144. I) Schematic explaining that proteasome inhibition should enhance ubi-sites actively targeting proteins for proteasomal degradation. J) As in H), for sites in ubi-site domain hotspots. The number of ubi-sites in each group from left-to-right are 23,758 and 503.

To understand the systems-level organisation of the ubiquitinome and its response to these perturbations, we first correlated and clustered the perturbation conditions (Fig. 3d). We generally observed medium-to-high correlations within the same perturbation across separate studies (Fig. S4a, average Pearson’s r = 0.498 between studies and r = 0.605 within studies). This highlights the reproducibility of the assembled data, especially considering studies did not necessarily employ the same treatment dose, treatment duration, cell line, or mass spectrometer.

Correlations in the response to different perturbations indicate potential functional relationships, such as the positive association between DNA damage-inducing UV radiation, protein folding stress, and ER stress (cluster III in Fig. 3d, example in Fig. 3e). These stressors all induce the unfolded protein response^73,74^, which triggers specific regulatory ubi-sites, including several sites on the ribosomal proteins RPS2 and RPS3 identified in our data^29^ (Fig. 3e). The unfolded protein response also globally inhibits protein synthesis^73^, leading to a widespread decrease of ubiquitination ordinarily resulting from the degradation of newly synthesised proteins^28^. In addition, the responses to UV and proteasome inhibition were negatively associated (Fig. 3d, f), particularly when the proteasome had already been inhibited before UV induction (Fig 3d, Fig. S4b). This may be explained by two mechanisms: first, degradative ubi-sites from newly synthesised proteins may decrease upon the UV-induced unfolded protein response but accumulate with acute proteasome inhibition^28^; second, this degradative chain buildup may limit ubiquitin availability for non-degradative DNA damage signals such as K6 and K63 chains^1,27,28,72^

We next assessed whether distinct patterns of ubi-site regulation are associated with differing levels of ubi-site conservation. Compared to all ubi-sites, highly conserved sites tended to be more up-regulated upon DNA damage, translation inhibition, and infection (Fig. 3g). In contrast, treatment with the proteasome inhibitors MG-132, bortezomib or epoxomicin enhanced the abundance of most poorly conserved sites, whereas many highly conserved sites failed to increase (Fig. 3g, h, Fig. S4c-d). Given that proteasome inhibition should cause the accumulation of ubi-sites actively targeting proteins for degradation (Fig. 3i), this observation implies that conserved ubi-sites are typically less likely to promote proteasomal degradation compared to the entire ubiquitinome. As an orthogonal measure of conservation we inspected our domain hotspots (Fig. 2f-j), and found that their ubi-sites were also less up-regulated upon proteasome inhibition (Fig. 3j, Fig. S4e-f). Taken together, these results suggest that ubi-sites that have been positionally conserved throughout evolution in order to support organismal fitness are less likely to promote proteasomal degradation, and instead may regulate protein function in signaling contexts such as the DNA damage response and infection.

### Prioritizing positionally important ubi-sites

Motivated by our analyses of ubi-site conservation and regulation, we sought a more general strategy to prioritise positionally important ubi-sites - site-specific ubiquitination events relevant for organismal fitness. Mirroring our previous work on phosphorylation^32^, we compiled 16 features spanning evolutionary, proteomic, structural, and other lines of evidence, capturing both empirical indicators of positional importance (e.g. ubi-site conservation, reliability of detection, regulation under perturbations) and the potential to modulate protein function (e.g. presence in a protein domain or at a predicted protein-protein interface, Fig. 4a, Table S6). We then integrated these features into a classifier trained to discriminate the 213 non-degradative, experimentally characterised ubi-sites in PhosphositePlus for which we could calculate features (Table S7, see Methods) from the 105,943 sites without functional annotation (“unannotated sites”). This approach rests on the assumption that the non-degradative regulatory sites are enriched for fitness-relevant, site-specific ubiquitination events while the unannotated sites are depleted in these, which is supported by our conservation analysis. The resulting classifier output can be interpreted as a “positional importance score” that prioritises known site-specific regulatory events; any unannotated sites that also score highly have putative positional importance.

**Figure 4:**
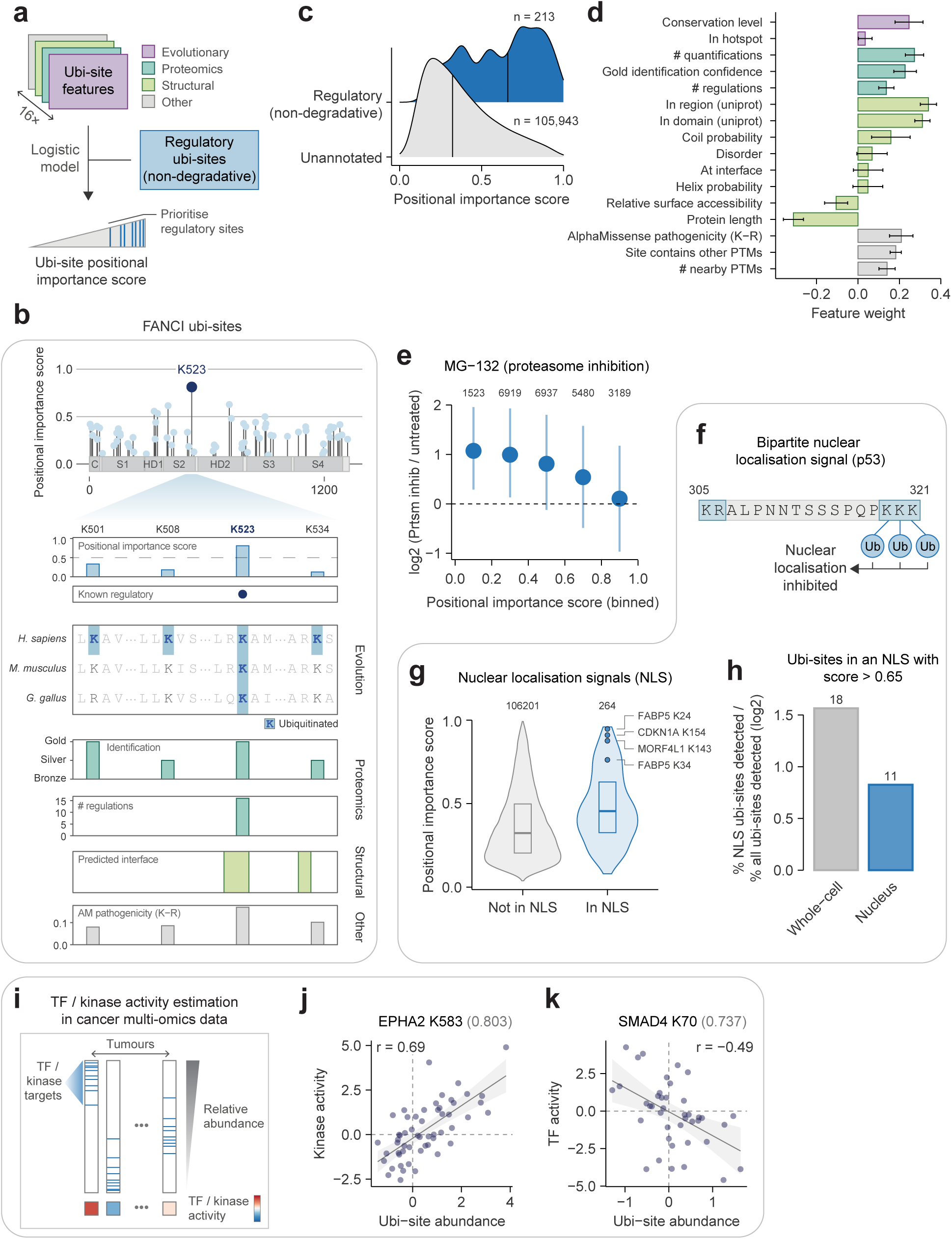
Prioritizing positionally important ubi-sites A) Overview of the workflow for generating a ubi-site positional importance score. Sixteen features capturing distinct biological dimensions were integrated into a logistic regression model trained to separate ubi-sites annotated for non-degradation regulatory functions from unannotated sites. B) Positional importance scores for ubi-sites on FANCI. The only regulatory site annotated in PhosphositePlus (K523) is colored dark blue. Select features used in the positional importance score are shown. C) Distribution of positional importance scores among ubi-sites annotated for non-degradation regulatory functions compared to unannotated ubi-sites. Numbers of unique ubi-sites per group are shown. D) Feature weights derived from logistic regression coefficients. Features with positive weights increase the positional importance score while those with negative weights decrease it. Error bars indicate standard deviation across multiple logistic regression models. E) Mean log2-fold changes (± s.d.) of ubi-sites under proteasome inhibition (MG-132 treatment) binned into positional importance score ranges. Numbers of quantified ubi-sites per bin are shown. Final values were obtained by normalising and averaging log2 fold-changes from multiple experiments (see Methods). F) Illustration of the p53 bi-partite nuclear localization signal (NLS), where ubiquitination inhibits nuclear localization^75^. G) Positional importance scores for ubi-sites within NLSs compared to other ubi-sites. Numbers of unique ubi-sites per group are indicated. Examples of high-scoring NLS-embedded ubi-sites are shown. H) The detection of NLS-embedded ubi-sites with positional importance score > 0.65 in a whole-cell proteome or nucleus-specific proteome from *Elia et al.*^27^, presented as numbers of unique sites (above barplot) and enrichment relative to all ubi-sites (barplot). I) Conceptual workflow for estimating kinase or transcription factor (TF) activity in tumors based on relative abundance of kinase/TF targets. J-K) Examples illustrating correlations between ubi-site abundance and J) kinase (EPHA2 K583) or K) TF (SMAD4 K70) activity across tumors. Pearson correlation coefficients (R), linear regression confidence intervals (95%), and ubi-site positional importance scores (plot titles) are shown.

Among several widely used classifiers - logistic regression, gradient boosting, and random forest - logistic regression demonstrated equivalent performance to the other more complex methods, achieving an average ROC AUC of 0.77 in five-fold train-test splits (Fig. S5a-b). Hence, we used logistic regression for final predictions. We then generated predictions by aggregating median scores from 25 logistic regression models trained through five time-repeated five-fold cross-validation, allowing us to establish robust scores (Table S6).

Our positional importance score successfully recovered well-known cases of site-specific regulatory ubiquitination, such as mono-ubiquitination of K523 on FANCI (Fig. 4b). This is the primary FANCI ubi-site coordinating FANCI/FANCD2-mediated interstrand crosslink repair^8^, and accordingly it received the highest positional importance score of all 67 ubi-sites identified in FANCI (Fig. 4b). Inspection of features used in the score reveals how they discriminate K523 from nearby ubi-sites less likely to be positionally important such as K501, K508, and K534; these sites are less conserved, less reliably identified, regulated in fewer proteomics experiments, and are not located at the FANCI-FANCD2 interface remodeled by K523 mono-ubiquitination (Fig. 4b).

On the other hand, many unannotated sites feature high scores equivalent to annotated regulatory sites, implying that a substantial fraction of the unexplored ubiquitinome may represent functionally important site-specific events (Fig. 4c). In particular, a score cutoff of 0.65 recovers approximately 50% of annotated non-degradative ubi-sites while classifying around 12,281 previously unannotated sites as positionally important. Logistic regression feature weights revealed that the score leveraged nearly all features to varying extents, with ubi-site presence within protein domains or regions, quantification in proteomics data, and conservation exerting the strongest positive influence (Fig. 4d). This underscores the value of our integrative machine-learning approach.

### Identifying regulatory roles of positionally important ubi-sites

We sought to investigate the molecular functions and regulatory properties captured by our ubi-site positional importance score. A recent landmark proteomics study measured ubi-site turnover and occupancy^34^; sites with a higher score exhibited faster turnover rates - consistent with dynamic regulatory roles - while no clear trend was observed for occupancy (Fig. S6a-b). Furthermore, lower-scoring sites were typically up-regulated upon proteasome inhibition by MG-132, bortezomib or epoxomicin, while higher-scoring sites showed limited regulation, suggesting they generally do not regulate protein function through proteasomal degradation (Fig. 4e, Fig. S6c-d).

To pinpoint the non-degradative functions played by high-scoring ubi-sites, we first asked whether they could affect protein localisation. Ubiquitination of nuclear localisation signals (NLSs) - which are characteristically rich in lysines - is likely to modulate nuclear import, exemplified by the bipartite NLS in p53^75^ (Fig. 4f). We identified 264 ubi-sites in nuclear localisation signals annotated in UniProt^76^; these NLS-embedded ubi-sites were enriched in higher positional importance scores, highlighting localisation modulation as a common outcome of positionally important ubi-sites. Furthermore, in a subcellular ubiquitin proteomics dataset we found that NLS-embedded ubi-sites with high positional importance scores (score > 0.65) were substantially less likely to be detected in the nuclear extract compared to whole-cell lysate^27^, providing evidence that ubiquitination of these sites may repress nuclear localisation (Fig. 4h).

Finally, we asked whether positionally important ubi-sites could modulate protein activity. To this effect, we estimated the activities of 44 protein kinases and 33 transcription factors (TFs) based on the relative abundance of known targets in phosphoproteomics and transcriptomics data from 85 lung squamous cell carcinoma tumours in the CPTAC consortium (Fig. 4i), which can be directly compared to ubiquitinomes measured on the same samples^77^. After controlling for the confounding effects of protein abundance and indirect regulation by protein degradation, we found six ubi-sites associated with the activity of their parent protein (see Methods, Table S8). The majority had high positional importance scores, implying that the score enriches for activity-regulating ubi-sites (Table S8). For instance, EPHA2 activity is positively associated with ubiquitination of K583 in the auto-inhibitory juxtamembrane domain^78^, which may function by relieving auto-inhibition (Fig. 4j). By contrast, activity of the TF SMAD4 is negatively associated with ubiquitination of K70 in the DNA-binding MH1 domain, which may interfere with DNA interactions (Fig. 4k). Overall, our positional importance score prioritises ubi-sites likely to regulate protein function through diverse avenues.

### Genetic characterization of high-scoring ubi-sites

Our positional importance score aims to identify site-specific ubiquitination relevant for organismal fitness. Next, we aimed to confirm this using clinical missense mutations from ClinVar, which can pinpoint protein residues critical to human health^79^. Mutations of ubiquitinated residues with higher positional importance scores were more likely to be pathogenic rather than benign or of uncertain effect, consistent with these sites performing critical biological functions (Fig. 5a). For instance, SMC1A K536 is located within the protein’s hinge domain that mediates dimerization with SMC3 in the Cohesin complex^80^, hence ubiquitination of this site may regulate complex assembly or crosstalk with other regulatory PTMs at this site^81^, and its disruption could underlie the cerebral congenital muscular hypertrophy associated with the K536R variant (Fig. 5b).

**Figure 5:**
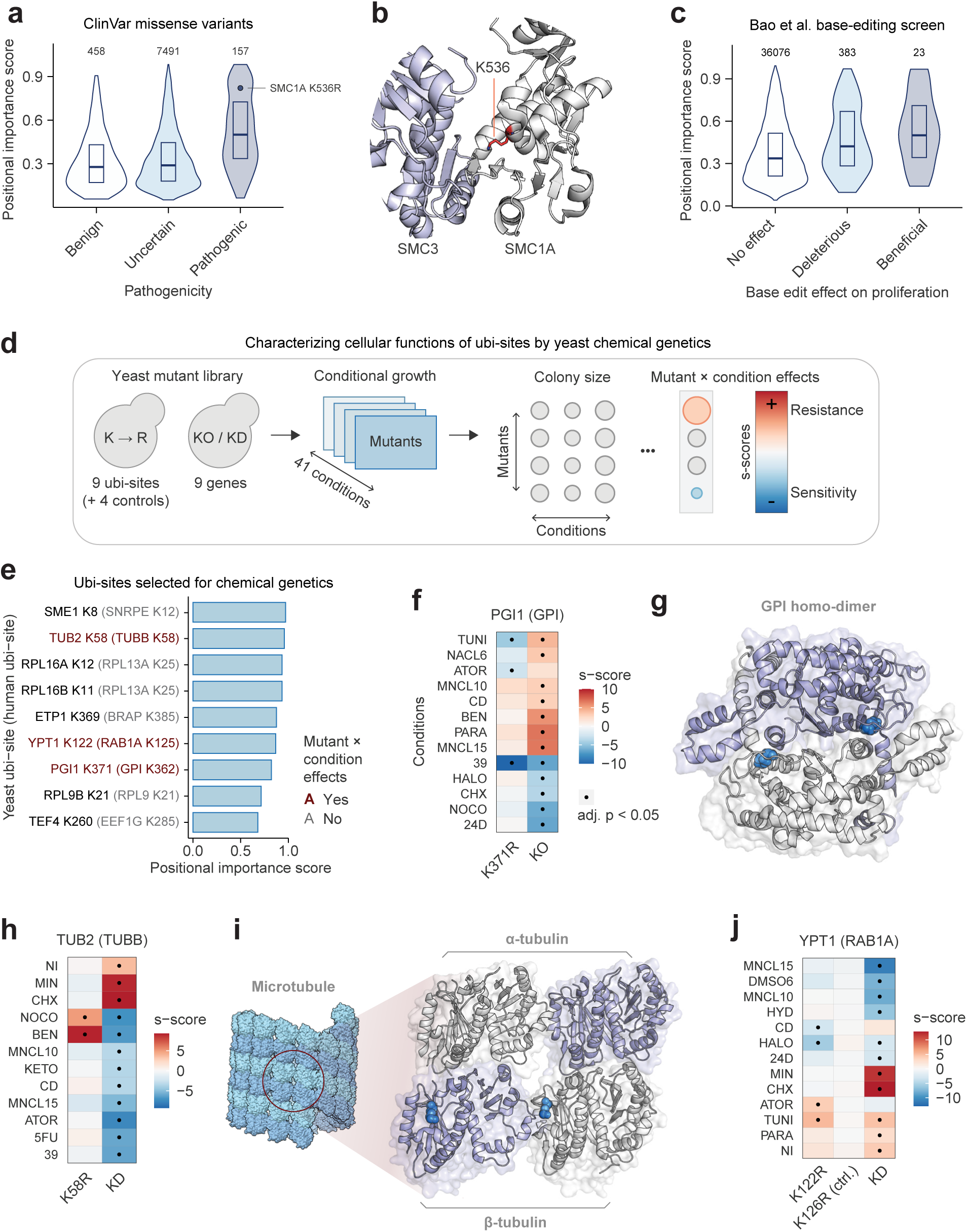
Genetic characterization of high-scoring ubi-sites A) Positional importance scores of ubi-sites co-localising with missense variants annotated in ClinVar as benign, uncertain, or pathogenic^79^. Numbers of unique ubi-sites per group are indicated. SMC1A K536R is shown. B) The interface of the SMC3 and SMC1A hinge domains within a Cryo-EM structure of the cohesin complex (PDB: 6WG4^80^). K536 on SMC1A is indicated in red. C) Positional importance scores of ubi-sites co-localising with lysine codon base edits in a base editing screen for hTERT-RPE1 cell proliferation^82^. D) Experimental workflow for characterizing yeast orthologs of human ubi-sites. Ubi-site lysine-to-arginine mutants, negative control mutants, and knockout/knockdown mutants of the corresponding genes were screened for conditional growth under 41 stress conditions (see Table S9 for descriptions of alleles and conditions). Growth phenotypes were quantified as s-scores, representing sensitivity or resistance under a given condition relative to other genotypes. E) Yeast ubi-sites (black text) and their human ubi-site orthologs (grey text in brackets) selected for experiments. Human ubi-site positional importance scores are indicated. Ubi-site mutants with stress-specific growth phenotypes are indicated in red. F) Significant growth phenotypes (adj. p < 0.05) for K371R or knock-out of yeast PGI1, corresponding to human GPI. G) Experimental structure of the human GPI homo-dimer (PDB: 9FHF^86^). The orthologous site of K371 is indicated in blue. H) Significant growth phenotypes (adj. p < 0.05) for K58R or knock-down of yeast TUB2, corresponding to human TUBB (β-tubulin). I) Left: Structural model of the microtubule with α-subunits in light-blue and β-subunits in dark-blue (PDB: 3J2U^87^). Right: Experimental structure of adjacent α-tubulin and β-tubulin subunits, with K58 indicated in blue (PDB: 6DPU^88^). J) Significant growth phenotypes (adj. p < 0.05) for K122R, K126R (a negative control), or knock-out of yeast YPT1, corresponding to human RAB1A.

To leverage more direct experimental evidence we consulted a published CRISPR-directed base-editing screen where 214,689 lysine codons were stochastically mutated to Arg, Glu, Gly, or Lys, identifying 1,572 lysine residues regulating hTERT-RPE1 cell proliferation^82^. Of the 36,873 lysine mutations occurring at ubi-sites, those affecting cell proliferation were enriched in higher positional importance scores, again supporting the importance of these positions for cellular homeostasis (Fig. 5c). To account for the possibility that the score enriches for proteins broadly susceptible to pathogenic variation at any residue, we directly compared ubi-sites with and without ClinVar/base-editing phenotypes within the same protein. Ubi-site scores were still higher for phenotypically active sites, although the differences were smaller than when examining the entire dataset, indicating some protein-level effects (avg. score difference = 0.094 ClinVar and 0.032 for base editing, Fig. S7a-b).

We reasoned that our positional importance should also highlight important regulatory positions across other species, due its enrichment for positional conservation. To this effect, we selected 8 high-scoring human ubi-sites from proteins performing diverse functions, and mutated the 9 orthologous yeast lysines to arginine using CRISPR-Cas9 (Fig. 5d-e, see Methods). We then screened site mutants for fitness effects across 41 stress conditions alongside knock-out or DaMP-mediated knock-down mutations^83^ of the corresponding gene, as well as negative control mutants of non-conserved lysines where possible, providing rich phenotypic information on ubi-site mutant effects (Fig. 5e, see Table S9 for descriptions of alleles and conditions).

Three of the nine ubi-site mutations impacted cell growth under one or more stress conditions (Fig. 5e). For both PGI1 K371 and TUB2 K58, most growth effects caused by ubi-site mutation were in the opposite direction to gene knockout/knockdown, consistent with ubi-site inhibitory functions (Fig. 5f-i). Ubiquitination may sterically hinder critical protein-protein interactions, since K371 lies at the interface of the enzymatically active PGI1 homodimer^84^ and K58 sits between adjacent TUB2 monomers within the microtubule (Fig. 5f-i). Alternatively, ubiquitination of these sites may trigger protein degradation. By contrast, mutating K122 in the GPTase YPT1 caused several fitness effects including resistance to the ER stress-inducing agent tunicamycin, consistent with YPT1’s role in combating ER stress through the unfolded protein response (Fig. 5j)^85^. Mechanistically, ubiquitination of K122 may affect nucleotide exchange, since this site lies in the conserved GTPase ubi-site hotspot discussed previously (Fig. 2i, S7c). Importantly, mutation of K126 - a nearby lysine not conserved in humans - exhibited no phenotype under any tested condition, confirming the specificity of our approach (Fig. 5j).

Ultimately, genetic evidence and mutagenesis experiments confirm that our positional importance score prioritizes ubi-sites maintaining cellular homeostasis through diverse regulatory mechanisms.

### Ubiquitination of ELAVL1 K320 disrupts RNA binding

We next aimed to leverage our positional importance score to uncover mechanistic insight into unstudied ubi-sites. We identified multiple high-scoring ubi-sites in RNA-binding protein and post-transcriptional regulator ELAVL1/HuR (Fig. 6a). K320 - the highest scoring ubi-site - resides at the interface between the third RNA recognition motif of ELAVL1 (RRM3) and 6×U RNA in a recently resolved crystal structure, suggesting that ubiquitination of K320 may modulate the interaction between ELAVL1 and RNA (Fig. 6a-b^89^). Supporting a non-degradational regulatory role, proteasome inhibition does not increase K320 ubiquitination (Fig. 6c). Furthermore, K320 ubiquitination is enhanced following UV exposure and during early Salmonella infection, implicating this site in ELAVL1’s known involvement in cellular stress and inflammation^90^ (Fig. 6c). These lines of evidence motivated further molecular dissection of K320 ubiquitination.

**Figure 6:**
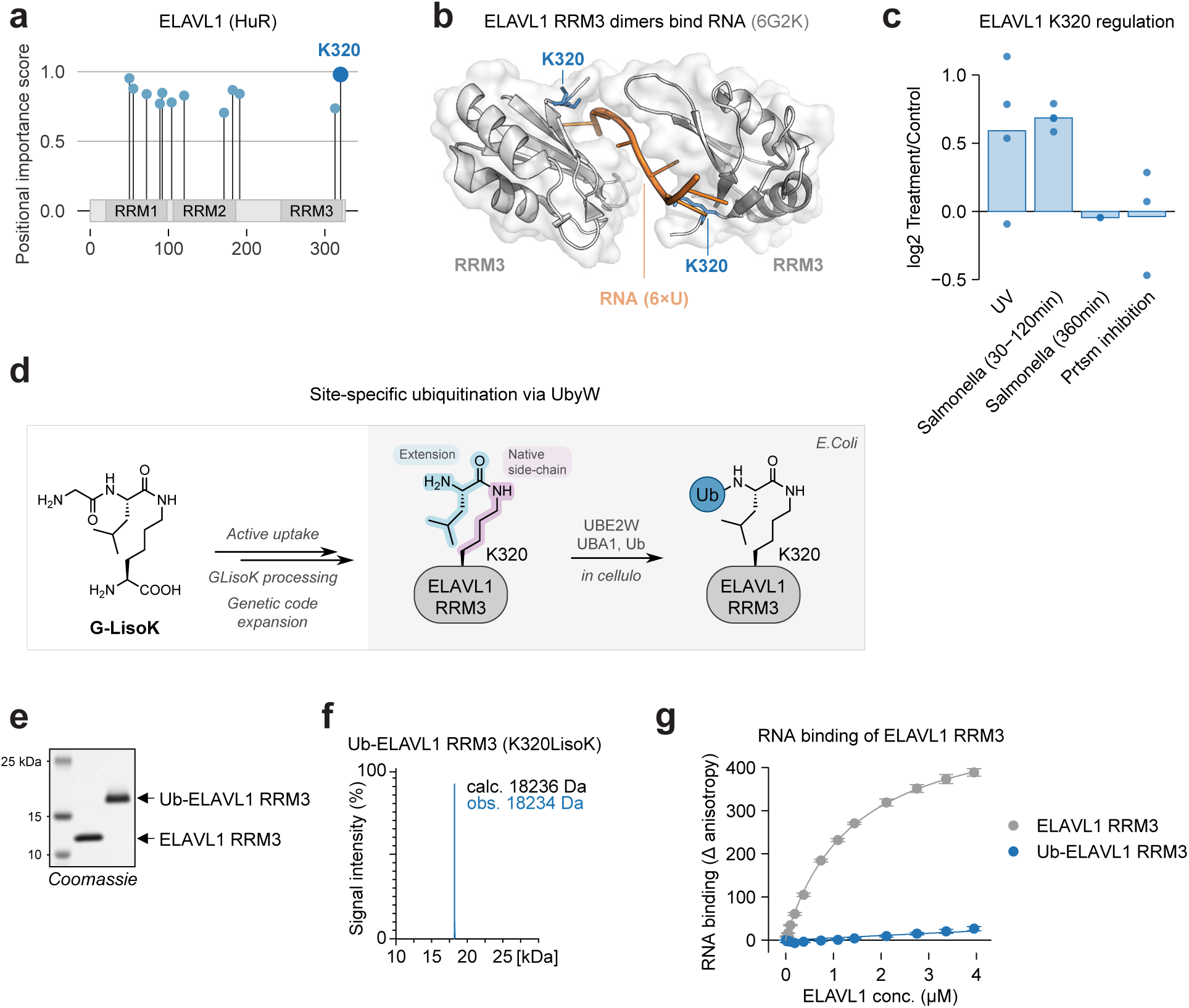
Ubiquitination of ELAVL1 K320 disrupts RNA binding A) Positional importance scores of ubi-sites on human ELAVL1 (HuR). B) Crystal structure of two ELAVL1 RRM3 domains binding 6×U RNA (6G2K^89^). C) Log2 fold changes of ELAVL1 K320 ubiquitination in ubiquitin proteomics data. D) Workflow for generating site-specifically ubiquitinated ELAVL1 RRM3 by UbyW^92^. E) Coomassie-stained SDS-PAGE of purified ELAVL1 constructs: Wildtype ELAVL1 RRM3 and ubiquitinated ELAVL1 RRM3 (K320LisoK). The full blot is shown in Fig. S8. F) LC-MS measurement of purified Ub-ELAVL1 RRM3. The non-deconvoluted spectrum is shown in Fig. S8. G) The binding of ELAVL1 RRM3 constructs to 10×U RNA was assessed by fluorescence anisotropy.

While it is straightforward to abolish ubiquitination by site mutagenesis, it is challenging to attach ubiquitin to a target lysine residue in a precise manner^91^. We employed our recently established workflow UbyW (Ubiquitylation by UBE2W)^92^ to site-specifically ubiquitylate the RRM3 domain of human ELAVL1 at position 320 (Fig. 6d). Briefly, we introduce the non-canonical amino acid LisoK at position 320 by amber suppression utilizing the active uptake of its propeptide precursor G-LisoK into *E. coli*^93^. Co-expressed UBE2W then recognizes the introduced neo-N-terminus of LisoK and conjugates ubiquitin onto it. This protein-ubiquitin conjugate recapitulates the native lysine-ubiquitin isopeptide bond with a single leucine extension at the C-terminus of ubiquitin (Fig. 6d).

Using this workflow we successfully generated and purified ubiquitinated ELAVL1 RRM3 (K320LisoK), as confirmed by SDS-PAGE and LC-MS (Fig. 6e-f, Fig. S8a-c). We then employed fluorescence anisotropy to assess binding to 10xU RNA, representing the U-rich sequences found in ARE-containing RNA^94^. Wildtype ELAVL1 RRM3 bound readily to 10xU RNA with a Kd of 1.4 μM, reflecting published results (Fig. 6g, Kd = 0.65 μM for the 11-mer RNA 5’-AUUUUUAUUUU-3’ in Ripin et al.^95^). By contrast, we observed virtually no RNA binding of ubiquitinated ELAVL1 RRM3 (K320LisoK), confirming our hypothesis that ubiquitination of this site modulates RNA binding (Fig. 6g). Overall, these results underscore the ability of our positional importance score to pinpoint regulatory ubiquitination events and guide elucidation of their molecular mechanisms.

## Discussion

Ubiquitination modulates protein function through diverse avenues in order to regulate a broad spectrum of critical cellular processes. However, 99% of catalogued human ubi-sites remain functionally uncharacterised^1,4,6,30^. Here, we have presented a functional analysis of the human ubiquitinome, comprising the generation of a high-confidence reference atlas of human ubi-sites and systems-wide analyses of ubi-site conservation and regulation, and culminating in a machine-learning score that prioritises site-specific regulatory events important for cellular homeostasis. We exemplified the utility of this score by exploring genetic vulnerabilities linked to ubiquitination, identifying ubi-sites involved in diverse regulatory mechanisms, and validating a subset of high-scoring sites experimentally.

We demonstrated that more evolutionarily conserved ubi-sites are less frequently linked to proteasomal degradation. This suggests that there is less selective pressure to maintain degradative ubiquitination at a specific location. Indeed, the mere presence of a degradative ubiquitin chain anywhere on a protein is generally sufficient for proteasome-mediated degradation, provided it is near a disordered initiation region that can be unraveled into the proteasome barrel^16^. On the other hand, the exact site of ubiquitination is likely often important for signaling functions including blocking or forming a protein-protein interaction interface^13–15^, altering protein conformations^8,96^, and modulating localisation signals^75^. The specific site of degradative ubiquitination may still matter in some cases, such as when only a subset of lysines are proximal to an initiation region^19^; when site-specific ubiquitination unfolds an otherwise ordered protein to provide an initiation region^17^; and when only a subset of lysines are targeted by a dedicated E3 ligase, such as those proximal to a phosphorylation-induced degron^20^. These cases may explain why we still observed highly conserved ubi-sites increasing in abundance upon proteasomal inhibition. Importantly, our results do not imply that protein degradation and quality control are unimportant for cellular homeostasis. Rather, there is likely functional redundancy at the level of the individual ubiquitination sites involved, rendering any single site less critical in isolation.

It is important to acknowledge some limitations of our results. Firstly, the annotated regulatory ubi-sites used to train the positional importance score may be shaped by biases, such as researcher preference to study ubi-sites that are conserved, detectable by mass spectrometry, and present in protein domains. Second, the di-glycine remnant antibody employed in most proteomics datasets in this study exhibits minor selectivity for surrounding amino acids^50^, fails to capture N-terminally linked ubiquitin, and also captures the ubiquitin-like modifications NEDD8 and ISG15 - estimated to represent 2% and 6% of the di-glycine remnant pool^28,52^. Future work will benefit from ubiquitin-specific enrichment methods such as the UbiSite protocol^26^. Third, in our pursuit of high-confidence ubi-site identifications, our reference ubiquitinome likely misses sites restricted to specific cell types or induced only under particular perturbations.

Another feature of ubiquitination we were unable to address is the chain architecture of ubiquitin modifications, which cannot be resolved at site resolution by current proteomics methods. Ubiquitin chain architectures confer distinct functional outcomes^1,4^, and it is possible that a given ubi-site performs different functions of differing importance depending on the chain type it carries. Recent advances in experimentally determining ubiquitin chain composition^97,98^ provide hope that our approach can eventually be extended to encompass ubi-site chain identity at proteome scale.

Overall, we have established a high-confidence reference ubiquitinome and provided a map of its evolutionary, regulatory, and functional terrain. By ranking approximately 100,000 human ubi-sites by their predicted positional importance, our resource opens new paths to a range of future mechanistic and functional studies across the community.

## Acknowledgements

First of all, we would like to thank all researchers who deposited the ubiquitin-enriched datasets in the PRIDE database. We would like to thank Steven Gygi and Woong Kim from Harvard Medical School for their assistance in generating proteomics data. J.A.V. would like to acknowledge funding from BBSRC [grant numbers BB/S01781X/1, BB/Y513829/1], EPSRC [grant number EP/Y035984/1], Wellcome Trust [grant number 223745/Z/21/Z] and EMBL core funding. A.J. acknowledges support from BBSRC grant [BB/S017054/1], and Doctoral Training Partnership (DTP) scholarship for EB [BB/T008695/1]. K.L. would like to acknowledge funding from ETH Zurich and the European Research Council (ERC under the European Union’s Horizon 2020 research and innovation programme, grant agreement no. 101003289-Ubl-tool to K.L). E.J.B acknowledges research funding provided by the NIH (R35GM148339, R01GM136994, DP2-GM119132).

## Author contributions

Conceptualization: P.B., I.B.H., E.J.B. Data curation: S.W., E.B., R.S., I.B.H. Formal analysis: J.vG., I.B.H., R.S., M.S., S.W., E.B. Funding acquisition: P.B., J.A.V., A.R.J., K.L., E.J.B. Investigation: M.F., B.B., P.S., A.C.C., M.A.B., E.R.T. Supervision: P.B., J.A.V., A.R.J., K.L., E.J.B., J.W.H., M.J.M. Visualization: J.vG. Writing – original draft: J.vG. Writing – review & editing: All authors

## Data availability

Proteomics data generated in this manuscript have been deposited to the ProteomeXchange Consortium via the PRIDE partner repository^33^ with the dataset identify PXD068906. The human reference ubiquitinome has been deposited to the ProteomeXchange with the dataset identify PXD068989.

## Code availability

All code used to analyse data and produce figures is available on request and will be made public before publication.

## Methods

### Computational analysis

Unless otherwise stated, computational analysis was performed using R (v4.3.1) in VSCode (v1.99.3). Visualisations were created using the ggplot2 package (v3.4.2).

### Creation of a human reference ubiquitinome

#### Selection of datasets

Datasets from the PRIDE database^33^ were selected for data re-analysis based on the following criteria: (i) human-derived samples enriched for lysine ubiquitination; (ii) data generated using Thermo Fisher Scientific instruments; and (iii) availability of metadata, either through the original publication or by direct communication with the authors.

After preliminary curation, 27 ubiquitination-enriched datasets were identified, of which 11 met the selection criteria, including employing different cell lines (e.g., HEK293, HeLa, HCT116). To ensure comparability, only datasets employing the diGly/UbiSite enrichment methods were included^24,26^. A summary of the resulting datasets, including PRIDE accession numbers, biological samples, and key characteristics, is provided in Table S1.

#### Proteomics raw data processing

Raw files from each dataset were converted to mzML format using ThermoRawFileParser (version 1.3.4) and analyzed independently. Dataset PXD037009 was split into 5 subsets because of its large size and the different experimental conditions included. An initial subset of raw files from each dataset was processed using Fragpipe with an open search to identify modifications. Modifications detected in > 1% of the peptide-spectrum matches (PSMs) were retained as parameters of the search. Peptide and protein identification, including post-translational modifications (PTMs), was performed using the Comet search engine (version 2024) on a Linux-based high-performance computing cluster. Default parameters were applied, with the following exceptions: missed cleavages were set to 4, and variable modifications were set to include PTMs from the Fragpipe open search results, with oxidation of methionine and N-terminal protein acetylation (excluding N-terminal peptide acetylation) included for all datasets. For all datasets, trypsin was used as a setting for digestion. The search database consisted of the UniProt human reference proteome (one protein per gene, downloaded in April 2024) and cRAP protein sequences as contaminants database (https://www.thegpm.org/crap/, obtained April 2024). Decoys were generated using the reverse decoy method via FragPipe (https://fragpipe.nesvilab.org/).

Statistical validation of PSMs and distinct peptide sequences was conducted using PeptideProphet and iProphet from the Trans-Proteomic Pipeline (TPP, version 7.1.0^99^).

High-confidence PSM matches were obtained, and PTM site localization was computed using PTMProphet (TPP), generating a unified mzidentML format file.

#### Post-processing

The searching result files from the TPP were processed using a custom Python script (mzidFLR; https://github.com/PGB-LIV/mzidFLR), as previously described^100^ and also applied in prior PTMeXchange projects^101^. First, a global false discovery rate (FDR) was calculated at the PSM level at a 1% threshold to retain high-confidence matches. The results were converted to a site-based format, with PTM localization scores assigned to each ubiquitination site. Contaminants, decoy hits, and non-ubiquitinated PSMs were excluded, retaining only ubiquitinated sites for downstream analysis.

Second, to estimate the probability of correct PTM localization, the PTM localization probability (from PTMProphet) was multiplied by the PSM probability (from PeptideProphet). Redundancy in PSMs site-based evidence was addressed by collapsing the data to the peptidoform level using a binomial adjustment^43^: the probability of a site being ubiquitinated was calculated by comparing the number of ubiquitination events to the total evidence for that site across all PSMs. False localization rates (FLRs) were estimated by introducing decoy alanine residues, yielding a 1% false discovery rate (FDR) PSM file and a PTM site-specific FLR file for quality control (QC) analysis for each individual dataset. In the case where a data set was analyzed in experimental blocks, the results were collapsed by retaining the evidence with the lowest FLR for a peptidoform. Peptide C-terminal ubiquitinated Lys were removed since it is expected that trypsin cannot cleave those. To combine results across datasets, a meta-analysis approach was employed to control FLR inflation. PTM sites were classified into Gold, Silver, and Bronze categories, with thresholds set as follows: Gold (≥ 2 datasets with FLR<1%), Silver (1 dataset with FLR<1%) and Bronze (≥ 1 datasets with FLR<5% and no datasets with FLR < 1%). When a peptide could be mapped to multiple protein sequences, the first protein in alphabetical order was selected to avoid redundancy.

#### Data set deposition

To ensure that the reanalyzed data meets FAIR guidelines^46^, the outputs have been made available in PRIDE with the identifier PXD068989.

### GO enrichment of highly ubiquitinated proteins

Gene ontology (GO) terms were extracted using UniProt protein accessions and the R packages org.Hs.eg.db (v3.17.0) and annotationDBI (v1.62.2). GO term enrichment was performed by one-side Fisher’s exact test on proteins with more than 10 ubi-sites, with the background as all proteins with at least one ubi-site. Only GO terms containing at least three process-regulating ubi-sites were tested. P-values were adjusted for within each ontology by the Benjamini-Hochberg procedure.

### PhosphositePlus annotations of regulatory ubi-sites

PhosphositePlus annotations of regulatory ubi-sites were downloaded (25 November 2024^30^). Ubiquitin site positions were corrected by manually aligned surrounding sequences into the respective protein sequence in the UniProt human proteome (19 February 2025). The “ON FUNCTION” column was used to assign proteins to degradative functions (“protein degradation”) or other functions.

### Ubiquitin proteomics experiments

For unpublished proteomics datasets, experiments were performed as described in Table S3. Mammalian cells were cultured as previously described^72^. Preparation of cell extracts for diGly-peptide immunoaffinity enrichment and MS/MS analysis were performed as previously described^28,72^. DiGly-peptides were quantified either by SILAC or label-free quantification as indicated in Table S3.

### Ubi-site conservation analysis

Protein multiple sequence alignments were extracted from EnsemblCompara GeneTrees^102^ (release 86) and subsetted for *H. sapiens, M. musculus, R. norvegicus, E. caballus, G. gallus, X. tropicalis, Danio rerio, D. melanogaster, C. elegans,* and *S. cerevisiae*. Ubi-site identifications from *H. sapiens, M. musculus, R. norvegicus, G. gallus, D. melanogaster, C. elegans,* and *S. cerevisiae* were mapped onto multiple sequence alignments. The lysine conservation at each alignment position was defined as the percentage of non-human residues that were lysine. The conservation level of each human ubi-site was scored using the following criteria:

Level 1: No ubiquitination in species apart from *H. sapiens*, lysine conservation < 10%.

Level 2: No ubiquitination in species apart from *H. sapiens*, lysine conservation > 10% and lysine conservation ≤ 50%.

Level 3: No ubiquitination in species apart from *H. sapiens*, lysine conservation > 50%.

Level 4: Ubiquitination detected in only two groups of species: 1) *H. sapiens* and 2) either *M. musculus or R. norvegicus*. Any lysine conservation level was allowed.

Level 5: Ubiquitination detected in three groups of species: 1) *H. sapiens*, 2) *M. musculus or R. norvegicus*, and 3) *G. gallus, D. melanogaster, C. elegans,* or *S. cerevisiae*. Any lysine conservation level was allowed.

### Ubi-site hotspot analysis

Ubi-site hotspots were identified using our pan-species ubiquitin proteomics data, including sites from *A. thaliana* that were not used in conservation analysis (Table S3 and Table S4). First, InterPro protein domains were extracted from UniProt protein identifiers using an API call (https://www.ebi.ac.uk/interpro/api/entry/interpro/protein/uniprot). Results were filtered for domains with a Pfam domain identifier. All domain sequences containing at least one ubi-site in any species were extracted. Domains with at least 15 instances and 50 ubi-sites were taken forward for further analysis. Domain sequences were aligned using MAFFT (v7.526, using the G-INS-i option^103^).

Hotspot analysis was then performed using domain alignments and ubi-site identifications as previously described^49^. Briefly, we first counted the observed ubi-sites at each position in a given domain alignment using a rolling window with a fixed size of 5 positions. To generate a null background model we randomly selected lysine residues within the alignment. Permutations were repeated 100 times and for each position in the alignment an expected median and standard deviation of phosphorylation were calculated. The observed values were converted to *z*-scores using the permutation information and then to *p*-values using the survival function of the normal distribution. Only enrichment over random was considered, and Bonferroni correction was used to account for multiple testing globally. To avoid identification of hotspots with a low effect size, a cut-off of an average of 2 ubi-sites per position was used. Finally, contiguous positions were merged to identify domain regions of interest with added ±2 positions on either side that were defined as phosphorylation hotspot regions. For each domain with a significant hotspot, the InterPro representative PDB structure was extracted/downloaded. The corresponding domain sequence was aligned to all other domain sequences using MAFFT (v7.526, using the G-INS-i option^103^). Hotspot analysis results were then mapped onto the representative sequence for visualisation. Hotspot analysis was performed with python v3.13.2.

### Quantitative ubiquitin proteomics data

#### Data preparation

Public proteomics data was accessed from the appropriate publications. Within each experimental condition log2 fold changes were calculated and averaged across different peptides corresponding to the same ubi-site, and across technical and biological replicates. Conditions with fewer than 500 quantified ubi-sites were filtered out.

#### Data analysis

Pairwise correlations between perturbation conditions were calculated using ubi-site log2-fold changes and Pearson’s correlation. The correlation heatmap was visualised using the R package ComplexHeatmap^104^ (2.16.0). Hierarchical clustering was performed on the heatmap using Pearson’s correlation as the distance metric. The clustering dendrogram was cut to create four clusters.

Log2-fold changes from different proteasome inhibitor experiments were combined to increase coverage. Specifically, within each proteasome inhibitor compound (MG-312, Bortezomib, and Epoxomicin) conditions were normalized by subtracting the condition median, dividing by the condition interquartile range, multiplying by the median of condition interquartile ranges, and adding the median of condition medians. Normalized log2-fold changes were then averaged across conditions, ignoring missing values.

### Machine learning for ubi-site positional importance scores

#### Feature collection

##### Conservation level

Conservation levels were converted into integers (1-5). Missing values from unaligned sites were imputed as the overall mean.

##### In hotspot

The presence of sites in ubi-site hotspots was encoded as a one-hot.

##### Gold identification confidence

Sites with Gold identification confidence were labeled with a one-hot encoding.

**# regulations**: Ubi-sites were defined as regulated in a given ubiquitin proteomics condition if they occurred in the top 5% or bottom 5% of log2-fold change values. The number of regulated conditions was counted at the site level, excluding proteasome inhibition conditions due to their distinct regulatory nature.

**# quantifications**: The number of quantified conditions was counted at the site level, including proteasome inhibition conditions.

**In domain, In region**: Protein domains and regions were extracted from UniProt using the GetFamily_Domains function from the R package UniprotR (2.3.0). Ubi-sites present in domains or regions were labeled with one-hot encodings.

**Disorder, helix probability, coil probability, relative surface accessibility**: NetSurfP3.0 was run on protein sequences to predict structural properties (v3.0^105^). Sequences larger than 5,000 residues could not be run and were hence excluded.

**Protein length**: Protein sequence length was calculated using canonical UniProt protein sequences.

**At interface**: Ubi-sites present at computationally predicted interfaces for 108,931 interacting protein pairs were labeled with a one-hot encoding (DockQ > 0.23^106^).

**Site contains other PTMs**: Ubiquitinated lysines annotated to contain other PTMs in PhosphositePlus (25 November 2024^30^) were labeled with a one-hot encoding.

**# nearby PTMs**: The number of PTMs recorded in (25 November 2024^30^) in a 21-residue window around each ubi-site was counted.

**AlphaMissense pathogenicity**: Pathogenicity scores were calculated for lysine to arginine mutations of each ubi-site using AlphaMissense^107^.

#### Feature processing

Several features were log-transformed to achieve more gaussian-like distributions (AlphaMissense pathogenicity, # nearby PTMs, # regulations, # quantifications, protein length). The subset of these features containing 0-values were transformed using the log(1 + x) function to avoid producing missing values. All features were centered and scaled using the preProcess function from the R package caret (version 6.0.94, method = c(“center”, “scale”)). Apart from ubi-site conservation levels, no missing values were imputed. Ubi-sites with missing values for any other feature were removed.

#### Model training and evaluation

The positive class was defined as all sites annotated to perform non-degradation functions in PhosphositePlus, and the negative class as all sites without an annotated function. The protein XRCC5 (UniProt: P13010) was removed from training data to mitigate bias. In particular, this protein contained 19 ubi-sites annotated to regulate DNA repair, which was by far the most number of regulatory sites per protein. Given the number of positive cases (n = 213), this risks overfitting to general properties of XRCC5 and its ubi-sites. Furthermore, the evidence underlying these annotations comes from experiments on a protein construct harbouring lysine-to-arginine mutations of all 19 sites at once^108^, which is weak evidence for the functional relevance of any one site in isolation.

Models were trained using the caret package in R. The model architectures used and their corresponding arguments for the caret “train” function were logistic regression (method = “glm”, family = “binomial”), gradient-boosting machine (method “gbm”) and random forest (method = “ranger”). In order to estimate the generalization error of these different architectures, models were trained and evaluated through train-test splits. Specifically, the data were divided into five equal folds; in each iteration, one fold was held out as the test set while the remaining four (80%) were used for training, yielding five distinct train–test splits. The createFolds function from caret was used to preserve the class balance. To prevent data leakage due to the use of protein-level features, ubi-sites belonging to the same protein were grouped together during fold separation. During each of the five train-test iterations, models were trained on the training data using a subsampling approach. In particular, five random samples were drawn from the negative class at 10× the size of the positive class, to mitigate the impact of the class imbalance and to speed up training. Within each subsample, all positive class instances were retained, and a model was training with 5-fold cross-validation for hyperparameter optimization. The optimized model was then applied to the corresponding test set, and model accuracy was measured using the area under the ROC curve (ROC AUC). The generalization error of each classifier architecture was then estimated by aggregating the 25 ROC AUC values arising from five negative subsamples within 5 train-test splits.

#### Generating predictions

To generate predictions, 25 train-test splits were generated by repeating the train-test split procedure described above five times. A logistic regression model was trained on each training set. Negative class subsampling was not used due to the speed of training logistic regression models; instead all samples were weighted inversely proportional to the size of the positive/negative class in order to mitigate the impact of the class imbalance. Each model was applied to all ubi-sites, generating prediction probabilities for the positive class. The median probability was calculated across the 25 models for each site to generate final positional importance scores.

### Transcription factor and kinase activity estimation from CPTAC data

Log2-transformed and normalized ubiquitin proteomics data were downloaded from the original publication (Table S3L^77^). The remaining CPTAC data were extracted using the cptac python package (cptac v1.5.14, python v3.12.4). The following pipelines were used: phosphoproteomics - bcm; proteomics - bcm; transcriptomics - bcm; whole-exome sequencing - harmonized. Non-tumour samples were removed. For phosphoproteomics, proteomics, and transcriptomics, data were transformed into log2 fold changes by subtracting the median within each tumour type for each quantified phosphosite/protein/transcript.

Transcription factor targets were extracted from the DoRothEa database (R package dorothea^109^ v1.12.0, confidence scores A-C). Kinase substrates were downloaded from PhosphositePlus (25 November 2024^30^). Kinase and transcription factor activities were estimated using the VIPER algorithm (viper() function from the R package viper^110^ v1.34.0) with minsize = 5, adaptive.size = FALSE, and eset.filter = F.

Ubi-sites associated with the activity of their parent protein were identified, applying a stringent multi-step procedure to exclude associations confounded by protein abundance and sites possibly regulating activity indirectly through protein degradation. In particular, the following steps were applied:

1. Retain ubi-sites significantly correlated with parent protein activity (Pearson’s correlation, Benjamini-hochberg adjusted p-value < 0.05).
2. Remove ubi-sites up-regulated upon proteasome inhibition (mean log2FC under Bortezomib and MG-132 treatment > 0),
3. For ubi-sites positively correlated with parent protein activity, regress protein abundance out of ubi-site abundance and protein activity using a linear model (“lm” function from the R package “stats” (version 4.5.1)). Repeat correlation between ubi-site and protein activity. Remove ubi-sites with unadjusted p-value ≥ 0.05.

### Nuclear localisation sequences

Nuclear localisation signals were extracted from UniProt using the GetFamily_Domains function from the R package UniprotR (2.3.0).

### Yeast chemical genomics

#### Overview of lysine point-mutant generation

Lysine point-mutant strains were generated using CRISPR-Cas9 in the yeast strain DHY214^111^ (a kind gift from Lars M. Steinmetz). Two confirmed mutant strains were used per point mutation to mitigate the risk of an off-target mutation causing the phenotype of interest, improving the reliability of the screen results. Point mutants were generated following a protocol from the Ellis lab (https://benchling.com/pub/ellis-crispr-tools), described in more detail below.

#### sgRNA design and cloning

Guide RNAs (gRNA) were designed to target sequences within 10 bp of an NGG or CCN PAM site. gRNAs are described in table S9. Forward and reverse oligonucleotides for gRNAs were ordered, phosphorylated using T4 PNK at 37 °C for 1 h, and annealed by heat denaturation and gradual cooling. Annealed oligonucleotides were ligated into BsmBI-digested entry vector pWS082 (a kind gift from Tom Ellis) using T7 ligase. Constructs were transformed into *E. coli* NEB 10β, and plasmids were recovered from GFP-negative colonies and confirmed by PCR.

#### Donor DNA construction

Donor fragments with 50 bp homology arms flanking the predicted Cas9 cleavage site (3 bp upstream of the PAM) were generated as overlapping oligonucleotides (20 bp overlap). Donor DNA sequences are described in table S9. Donors were assembled by PCR with Phire DNA polymerase, followed by ethanol precipitation and elution.

#### Plasmid preparation

sgRNA plasmids were digested with EcoRV and pooled when multiple guides were used. Cas9-sgRNA gap repair plasmids (pWS173, a kind gift from Tom Ellis) were linearized with BsmBI, gel-purified, and adjusted to 100 ng/µl.

#### Yeast transformation

Yeast transformations were performed as previously described with some modifications^112^. First, DHY214 yeast cells were grown to exponential phase, harvested by centrifugation and washed with LiAc 100 mM, 1× TE pH 8.0 buffer to produce chemically competent yeast cells. DNA was added to cells (100 ng linearized Cas9-sgRNA gap repair plasmid, 200 ng digested sgRNA plasmid, and 3-5 µg donor DNA) followed by transformation mix: 100 µl of 50% PEG 3350, 15 µl of 1 M LiAc, 20 µl of single-stranded carrier DNA from salmon sperm and 30 µl of competent cells. The resulting mix was heat-shocked at 42 °C for 40 min with shaking at 600 rpm in a thermomixer. Outgrowth was performed in 1.5 ml tubes in YPAD medium at 30 °C, shaking at 1500 rpm on a thermomixer for ∼4 h, after which colonies were plated onto YPD + NAT (100 µg/mL Nourseothricin). 4-8 colonies were sequenced to select for successful editing.

#### Chemical genomics screening

The BY4741 *MATa* haploid KO library, DaMP knock-down library^83^, and the lysine point-mutant library were maintained in YPAD + G418, YPAD + G418 and YPAD, respectively, before screening in 384 colony format. All strains used in this study are described in Table S9 and were maintained on the appropriate selection media. chemical genomics screening was performed as previously described^112^. All chemical genomics growth conditions are described in Table S9. Colony imaging and calculation of s-scores and q-values were performed as previously described^112^. For downstream analysis and visualisation, s-scores and q-values were collapsed across replicates and timepoints by calculating the mean (s-scores) or geometric mean (q-values).

### ELAVL1 ubiquitination experiments

#### General Methods

Codon optimized genes encoding ELAVL(244-326)-K320TAG and SUMO2-ELAVL(244-326)-wt were purchased as DNA Strings (Twist Bioscience) and cloned into the pBAD vectors via restriction cloning. All solvents and chemical reagents were purchased from Sigma Aldrich, Senn, Carbolution, Acros Organics, or Fisher Scientific and were used without further purification unless stated otherwise. Bolt 4-12 % Bis-Tris gradient gels (Invitrogen) were run (at 165 V for 40 min) using a BoltTM Mini Gel Tank system (Invitrogen). Gels were stained with Quick Coomassie Stain (Generon). PageRuler Prestained Plus Protein Ladder 10-250 kDa (ThermoFisher) was used as the protein marker. Protein and DNA concentrations were measured on a NanoPhotometer® NP60 (Implen). Protein LC-MS was performed on an Agilent Technologies 1260 Infinity LC-MS system with a 6310 Quadrupole spectrometer equipped with a Phenomenex Jupiter C4 300 A LC Column (150 x 2 mm, 5 μm). The solvent system consisted of 0.1% formic acid in water (solvent A) and 0.1% formic acid in acetonitrile (solvent B). HPLC-purified 5’-FAM labeled 10xU RNA was obtained from Microsynth. Fluorescence anisotropy measurements were performed on a Jasco Fluorescence Spectrometer FP-8350 equipped with polarizers (Jasco). GLisoK was prepared as previously described^93^.

#### Expression and purification of ELAVL-wt

Chemically competent E. coli K12 cells were transformed with pBAD H6-SUMO-ELAVL-wt (encoding for H6-SUMO-SGSG-ELAVL(244-326) wt (see plasmids in Table S10 and methods section “Protein sequences”)). After recovery with 1 mL of SOC medium for 1 h at 37 °C, the cells were cultured overnight in 50 mL of 2× YT medium supplemented with ampicillin (100 µg/mL) at 37 °C, 200 rpm. The overnight culture was diluted to an OD600 of 0.05 in 200 mL of fresh autoinduction medium^113^ supplemented with ampicillin (50 µg/mL) and cultured at 37 °C, 200 rpm overnight. The cells were harvested by centrifugation (4000 × g, 20 min, 4 °C) and resuspended in 15 mL of lysis buffer (20 mM Tris pH 7.5, 300 mM NaCl, 30 mM imidazole, 0.175 mg/mL PMSF). The cell suspension was incubated on ice for 30 min and sonicated with cooling in an ice-water bath. The lysed cells were centrifuged (14,000 × g, 20 min, 4 °C), the cleared lysate added to Ni Sepharose 6 Fast Flow (Cytiva) (0.5 mL of slurry per 1 L of culture) and the mixture was incubated with agitation for 1 h at 4 °C. After incubation, the mixture was transferred to an empty plastic column and washed with 10 CV (column volumes) of wash buffer (20 mM Tris pH 7.5, 300 mM NaCl, 30 mM imidazole). On beads SUMO-cleavage was performed by addition of SUMO protease (Sigma-Aldrich, Cat. No. SAE0067) in wash buffer supplemented with 1 mM TCEP, followed by incubation at 4 °C for 3 h. The flow through was collected and applied to size-exclusion chromatography (SEC) using a Superdex Increase 75 10/300 (GE Healthcare) with SEC buffer (20 mM Tris pH 7.5, 100 mM NaCl and 1 mM DTT). Fractions containing the ELAVL wt were pooled together and concentrated using Amicon centrifugal filter units with a 3 kDa MWCO (Millipore). Protein concentration was calculated from the measured A280 absorption (extinction coefficients were calculated with ProtParam (https://web.expasy.org/protparam/)).

#### Expression and purification of site-specifically ubiquitylated Ub-ELAVL(K320LisoK) via a UBE2W-based reconstituted ubiquitylation cascade in E. coli

Chemically competent E. coli K12 cells were cotransformed with pBAD H6-TEV-ELAVL K320TAG / UBE2W (encoding for H6-TEV-GS-ELAVL(244-326)-K320TAG and UBE2W (D. rerio)) and pEVOL Ub/E1/Ma aaRS/tRNA (encoding for ubiquitin, UBA1, Ma “IP” aaRS and Ma tRNA (see plasmids in Table S10 and methods section “Protein sequences”)). After recovery with 1 mL of SOC medium for 1 h at 37 °C, the cells were cultured overnight in 50 mL of 2× YT medium supplemented with ampicillin (100 µg/mL) and chloramphenicol (50 µg/mL) at 37 °C, 200 rpm. The overnight culture was diluted to an OD600 of 0.05 in 200 mL of fresh autoinduction medium^113^ supplemented with ampicillin (50 µg/mL) and chloramphenicol (25 µg/mL) and GLisoK (500 µM) and cultured at 37 °C, 200 rpm overnight. The cells were harvested by centrifugation (4000 × g, 20 min, 4 °C) and resuspended in 15 mL of lysis buffer (20 mM Tris pH 7.5, 300 mM NaCl, 30 mM imidazole, 0.175 mg/mL PMSF). The cell suspension was incubated on ice for 30 min and sonicated with cooling in an ice-water bath. The lysed cells were centrifuged (14,000 × g, 20 min, 4 °C), the cleared lysate added to Ni Sepharose 6 Fast Flow (Cytiva) (0.5 mL of slurry per 1 L of culture) and the mixture was incubated with agitation for 1 h at 4 °C. After incubation, the mixture was transferred to an empty plastic column and washed with 10 CV (column volumes) of wash buffer (20 mM Tris pH 7.5, 300 mM NaCl, 30 mM imidazole) followed by elution in 150 µL fractions with wash buffer supplemented with 300 mM imidazole. Afterwards, TEV Protease (Sigma-Aldrich, Cat. No. T4455) was added to the elution and incubated at 4 °C for 3 h. TEV Protease was removed by reverse NiNTA purification followed by SEC using a Superdex Increase 75 10/300 (GE Healthcare) with a SEC buffer (20 mM Tris pH 7.5, 100 mM NaCl and 1 mM DTT) to separate ubiquitylated ELAVL from unmodified ELAVL. Fractions containing Ub-ELAVL(K320LisoK) were pooled together and concentrated using Amicon centrifugal filter units with a 10 kDa MWCO (Millipore). Protein concentration was calculated from the measured A280 absorption (extinction coefficients were calculated with ProtParam (https://web.expasy.org/protparam/)).

#### Kd determination via fluorescence anisotropy

Fluorescence anisotropy measurements were conducted on a Jasco Fluorescence Spectrometer FP-8350 equipped with polarizers (Jasco). Excitation and emission monochromators were set to 500 nm and 520 nm and measurements were performed at RT with a bandwidth of 5 nm. ELAVL/Ub-ELAVL(K320LisoK) were titrated to 100 nM 5’FAM labeled 10xU RNA in 20 mM Tris pH 7.5, 200 mM NaCl, 1 mM DTT and 0.01% NP40. All data processing was performed using GraphPad Prism 10 (GraphPad software). Kd was determined using a single binding site model, and average values and error bars (s.d.) were calculated from three different experiments (n = 3).

#### Protein Sequences

H6-SUMO-SGSG-ELAVL-wt MPGSHHHHHHGSDSEVNQEAKPEVKPEVKPETHINLKVSDGSSEIFFKIKKTTPLRRLMEAFAKRQGKEMDSLRFLYDGIRIQADQTPEDLDMEDNDIIEAHREQIGGSGSGWCIFIYNLGQDADEGILWQMFGPFGAVTNVKVIRDFNTNKCKGFGFVTMTNYEEAAMAIASLNGYRLGDKILQVSFKTNK SHK

H6-TEV-GS-ELAVL-K320TAG MPHHHHHHGENLYFQGSGWCIFIYNLGQDADEGILWQMFGPFGAVTNVKVIRDFNTNKCKGF GFVTMTNYEEAAMAIASLNGYRLGDKILQVSF*TNKSHK

Ubiquitin MQIFVKTLTGKTITLEVEPSDTIENVKAKIQDKEGIPPDQQRLIFAGKQLEDGRTLSDYNIQKESTL HLVLRLRGG

UBA1 MAKNGSEADIDEGLYSRQLYVLGHEAMKRLQTSSVLVSGLRGLGVEIAKNIILGGVKAVTLHDQGTAQWADLSSQFYLREEDIGKNRAEVSQPRLAELNSYVPVTAYTGPLVEDFLSGFQVVVLTNTPLEDQLRVGEFCHNRGIKLVVADTRGLFGQLFCDFGEEMILTDSNGEQPLSAMVSMVTKDNPGVVTCLDEARHGFESGDFVSFSEVQGMVELNGNQPMEIKVLGPYTFSICDTSNFSDYIRGGIVSQVKVPKKISFKSLVASLAEPDFVVTDFAKFSRPAQLHIGFQALHQFCAQHGRPPRPRNEEDAAELVALAQAVNARALPAVQQNNLDEDLIRKLAYVATGDLAPINAFIGGLAAQEVMKACSGKFMPIMQWLYFDALECLPVDKEVLTEDKCLQRQNRYDGQVAVFGSDLQEKLGKQKYFLVGAGAIGCELLKNFAMIGLGCGEGGEIIVTDMDTIEKSNLNRQFLFRPWDVTKLKSDTAAAAVRQMNPHIRVTSHQNRVGPDTERIYDDDFFQNLDGVANALDNVDARMYMDRRCVYYRKPLLESGTLGTKGNVQVVIPFLTESYSSSQDPPEKSIPICTLKNFPNAIEHTLQWARDEFEGLFKQPAENVNQYLTDPKFVERTLRLAGTQPLEVLEAVQRSLVLQRPQTWADCVTWACHHWHTQYSNNIRQLLHNFPPDQLTSSGAPFWSGPKRCPHPLTFDVNNPLHLDYVMAAANLFAQTYGLTGSQDRAAVATFLQSVQVPEFTPKSGVKIHVSDQELQSANASVDDSRLEELKATLPSPDKLPGFKMYPIDFEKDDDSNFHMDFIVAASNLRAENYDIPSADRHKSKLIAGKIIPAIATTTAAVVGLVCLELYKVVQGHRQLDSYKNGFLNLALPFFGFSEPLAAPRHQYYNQEWTLWDRFEVQGLQPNGEEMTLKQFLDYFKTEHKLEITMLSQGVSMLYSFFMPAAKLKERLDQPMTEIVSRVSKRKLGRHVRALVLELCCNDESGEDVEVPYVRYTIR Ma PylRS H227I Y288P

MMTVKYTDAQIQRLREYGNGTYEQKVFEDLASRDAAFSKEMSVASTDNEKKIKGMIANPSRHGLTQLMNDIADALVAEGFIEVRTPIFISKDALARMTITEDKPLFKQVFWIDEKRALRPMLAPNLYSVMRDLRDHTDGPVKIFEMGSCFRKESHSGMHLEEFTMLNLVDMGPRGDATEVLKNYISVVMKAAGLPDYDLVQEESDVYKETIDVEINGQEVCSAAVGPIPLDAAHDVHEPWSGAGFGLERLLTIREKYSTVKKGGASISYLNGAKIN

## Supplementary figures

**Figure S1:**
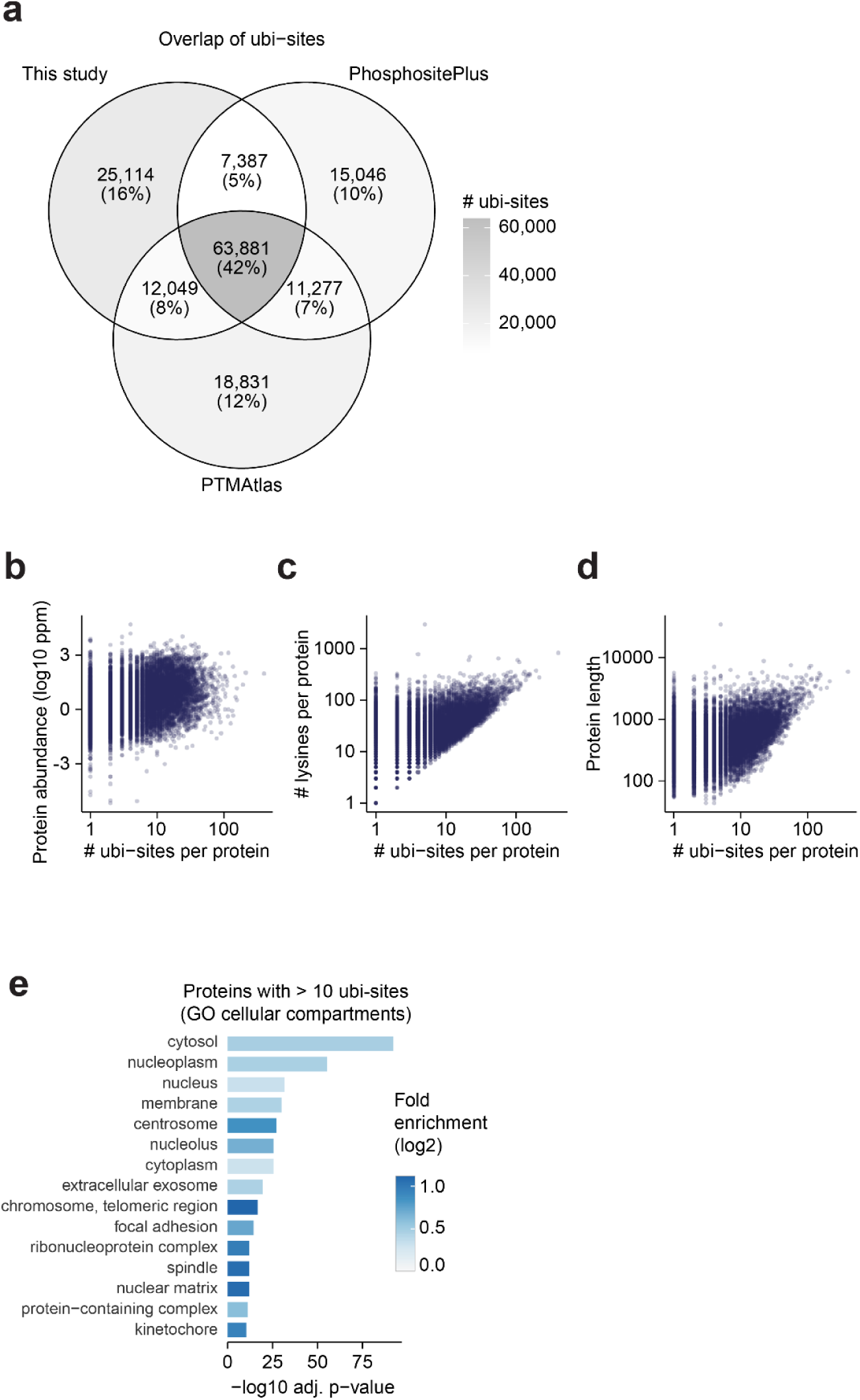
Additional analysis of reference ubiquitinome A) The overlap of unique ubi-sites identified in our reference ubiquitinome, PhosphositePlus^30^, and PTMAtlas^44^ is shown. B-D) The relationship betwen the number of ubi-sites identified per protein and B) protein abundance, C) protein length, and D) the number of lysines in a protein sequence. E) The top 15 significantly overrepresented GO cellular compartment terms among proteins with more than 10 ubi-sites, relative to all proteins in the human reference ubiquitinome (one-sided Fisher’s exact test, Benjamini-Hochberg p-value adjustment).

**Figure S2:**
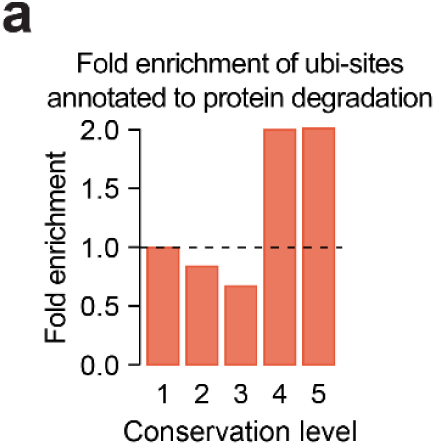
Additional analysis of conserved ubi-sites A) The enrichment of ubiquitin sites annotated to perform degradation regulatory functions at each conservation level (PhosphositePlus^30^).

**Figure S3:**
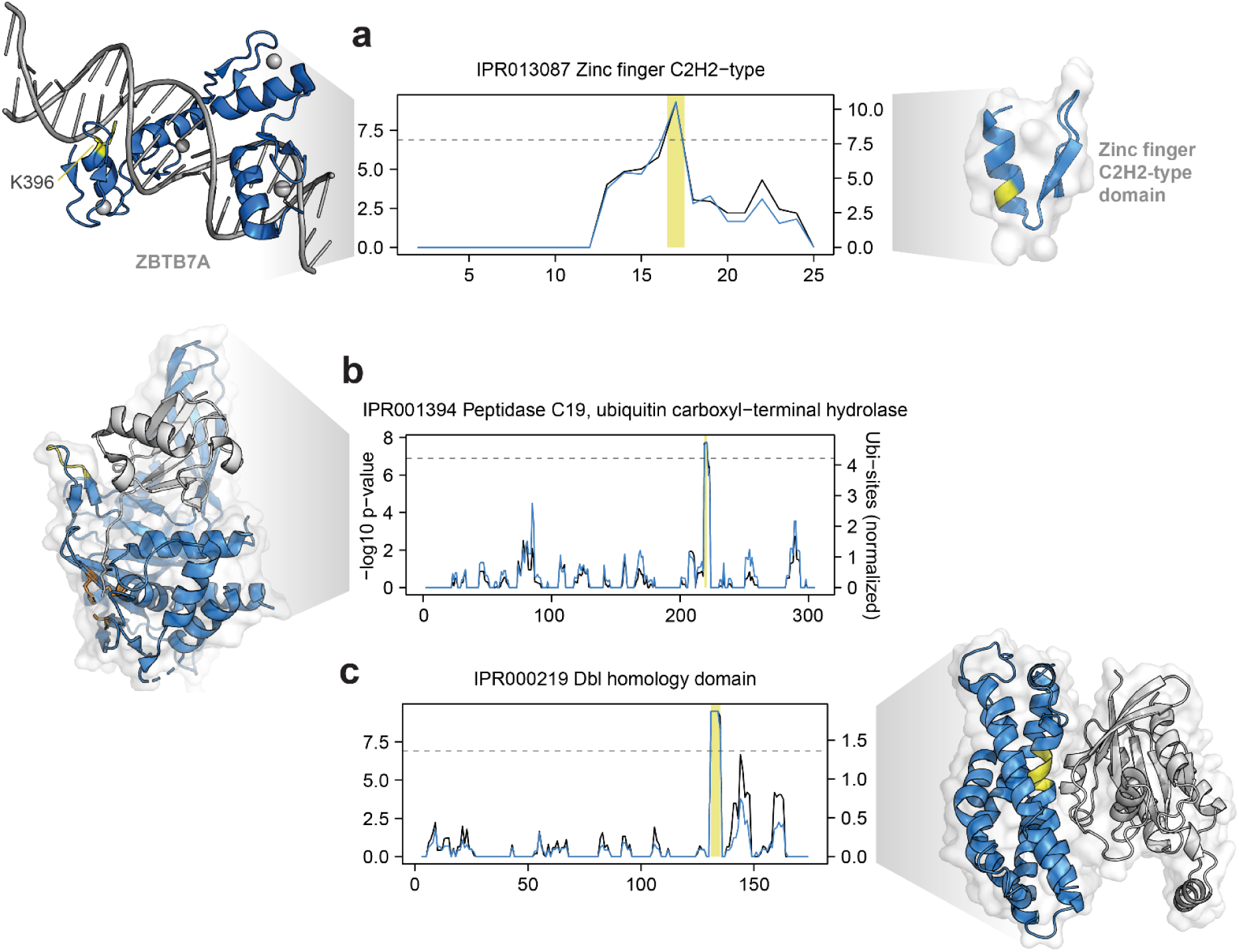
Additional ubi-site hotspots A-C) Identification of ubi-site hotspots in three protein domains. The black line indicates the average number of ubi-sites observed across the domain sequence alignment within a rolling window, normalised by subtracting the number of ubi-sites expected by chance. The blue line indicates the p-value associated with the enrichment of ubi-sites at each alignment position. The horizontal line indicates a Bonferroni-corrected p-value cut-off of 0.01 (uncorrected p-value < 1.29×10^-7^). Positions with a-log10 p-value above this cut-off and average number of phosphosites per window higher than 2 are classified as hotspot regions and highlighted with a yellow bar. Hotspot regions are mapped onto representative structures for each domain in yellow. In A) an example of a hotspot ubi-site located at the interface of a zinc finger with DNA is shown (K396 in ZBTB7A, PDB: 8E3E^114^). In B) the catalytic triad of the protease is coloured orange and a ubiquitin molecule covalently bound to the catalytic triad is coloured white. In C) the interaction partner Rac1 is coloured in white.

**Figure S4:**
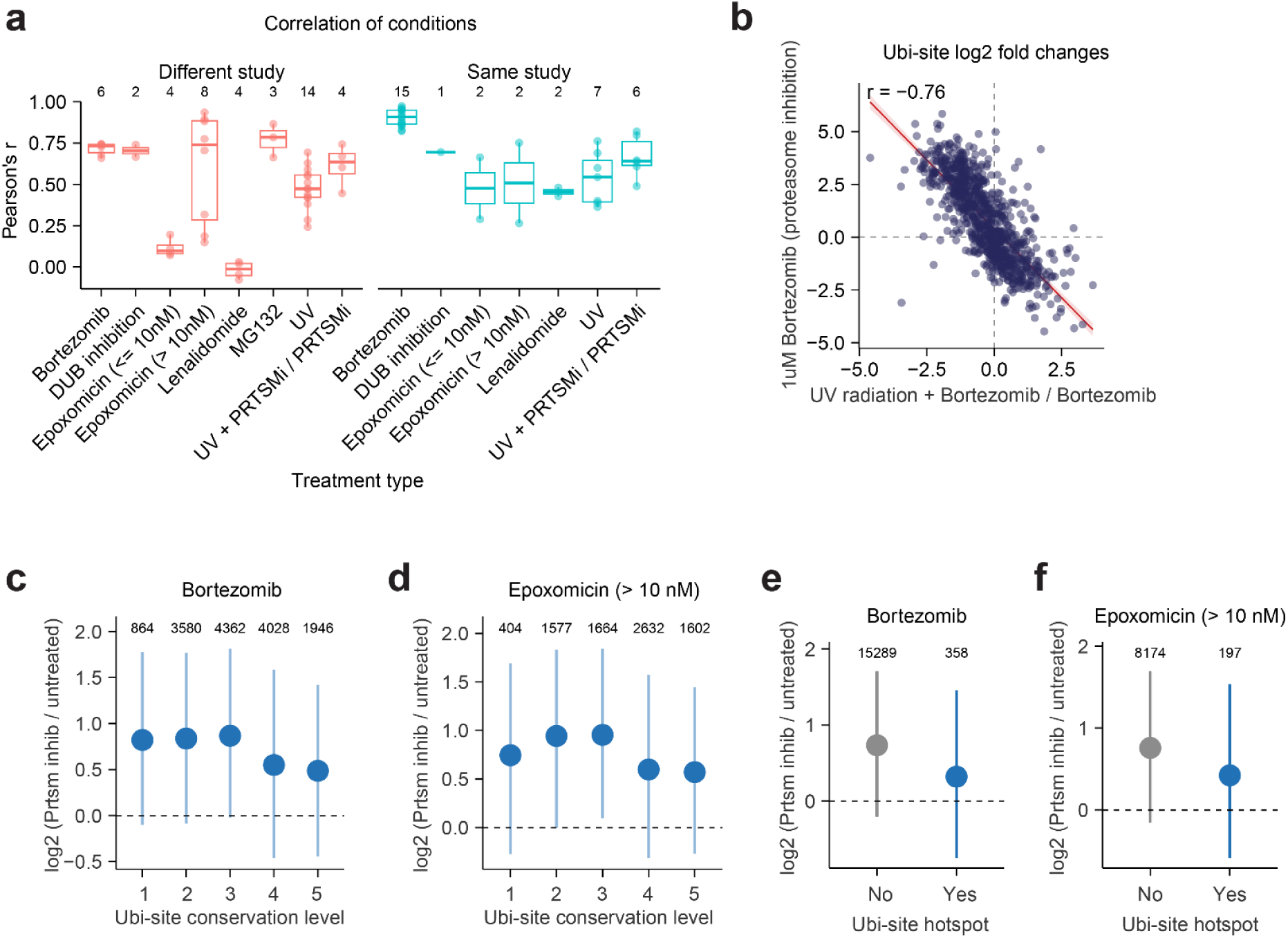
Additional analysis of quantitative ubiquitin proteomics data A) Pearson’s correlation was performed using ubi-site fold changes between pairs of conditions representing the same treatment. Pairs were separated based on whether the conditions came from the same study or different studies. The number of pairs in each group is indicated above boxplots. B) The correlation in ubi-site fold changes between Bortezomib treatment and UV radiation on a background of Bortezomib treatment. C-D) Mean log2-fold changes (± s.d.) of ubi-sites under proteasome inhibition (bortezomib or epoxomicin treatment), grouped by evolutionary conservation level. Final values were obtained by normalising and averaging log2 fold-changes from multiple experiments (see Methods). The number of ubi-sites in each group is indicated. Only epoxomicin treatments at > 10 nM were used since lower doses do not cause proteome-wide inhibition of protein degradation^72^. E-F) As in C-D), for sites in ubi-site domain hotspots.

**Figure S5:**
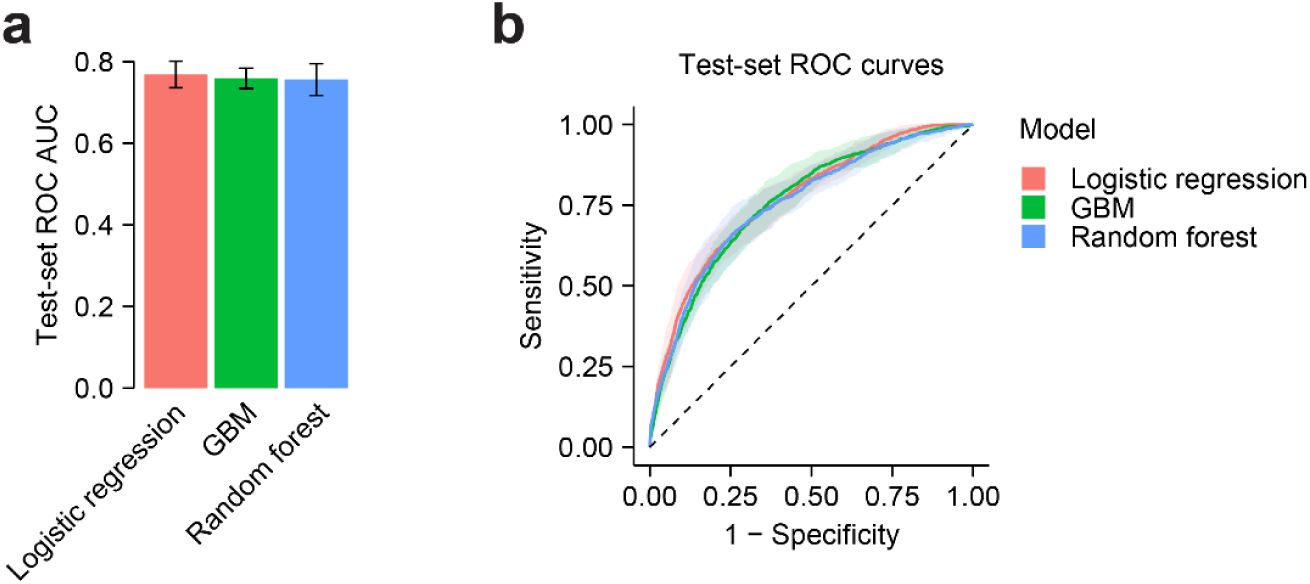
Training a machine-learning based ubiquitin site positional importance score A-B) Three classifier architectures were trained to separate unannotated ubi-sites from ubi-sites annotated to non-degradation roles. Classifiers were trained and evaluated through a five-fold train-test split, with hyperparameter optimization performed during each training iteration via five-fold cross-validation. A) shows ROC curve AUCs on each test set (error bars indicate standard deviation) while B) shows averaged ROC curves (shading indicates standard deviation).

**Figure S6:**
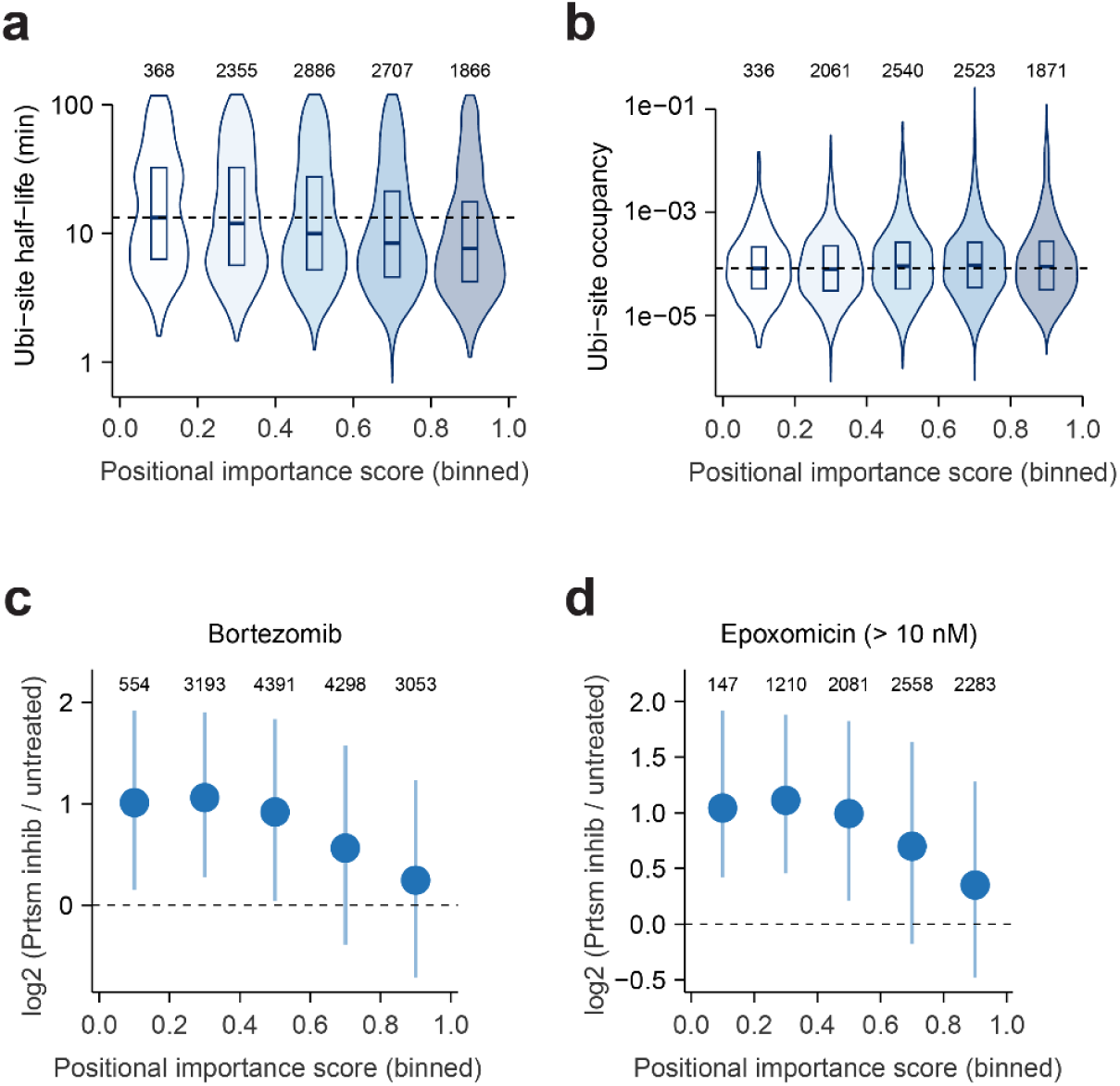
Additional characterisation of the ubi-site positional importance score A-B) Measurements of A) ubi-site half-life and B) ubi-site occupancy in *Prus et al*.^34^, binned into positional importance score ranges. Numbers of quantified ubi-sites per bin are shown. The median of the lowest positional importance score bin is shown as a dotted line. C-D) Mean log2-fold changes (± s.d.) of ubi-sites under proteasome inhibition (bortezomib or epoxomicin treatment), binned into positional importance score ranges. Numbers of quantified ubi-sites per bin are shown. Final values were obtained by normalising and averaging log2 fold-changes from multiple experiments (see Methods). Only epoxomicin treatments at > 10 nM were used since lower doses do not cause proteome-wide inhibition of protein degradation^72^.

**Figure S7:**
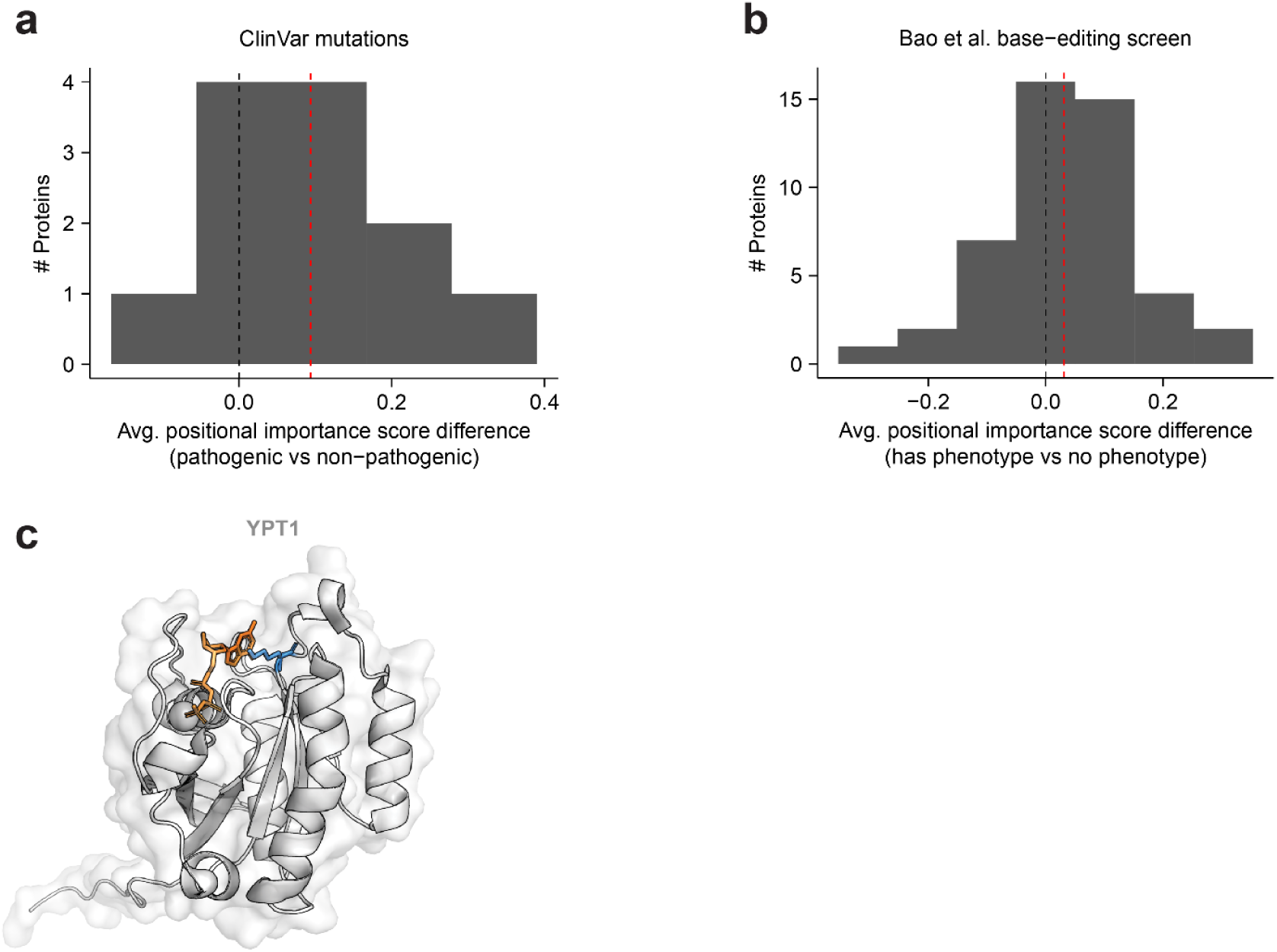
Additional genetic analysis of high-scoring ubi-sites A) ClinVar missense variants co-localising with ubi-sites were collected. Only proteins with at least two pathogenic and two non-pathogenic ClinVar ubi-site variants were examined for robustness. Within each protein, the average ubi-site positional importance score was compared between lysines with and without pathogenic variants. The red dotted line indicates the mean difference across proteins and the black dotted line indicates a value of 0 for comparison. B) The analysis in A) was repeated using base editing phenotypes from Bao et al.^82^. The comparison was performed between lysines with base edits that affected proliferation and those with base edits that didn’t affect proliferation. C) Experimental structure of yeast YPT1 (PDB: 1YZN^115^). K122 is indicated in blue and the GTP analog GppNHp in orange.

**Figure S8:**
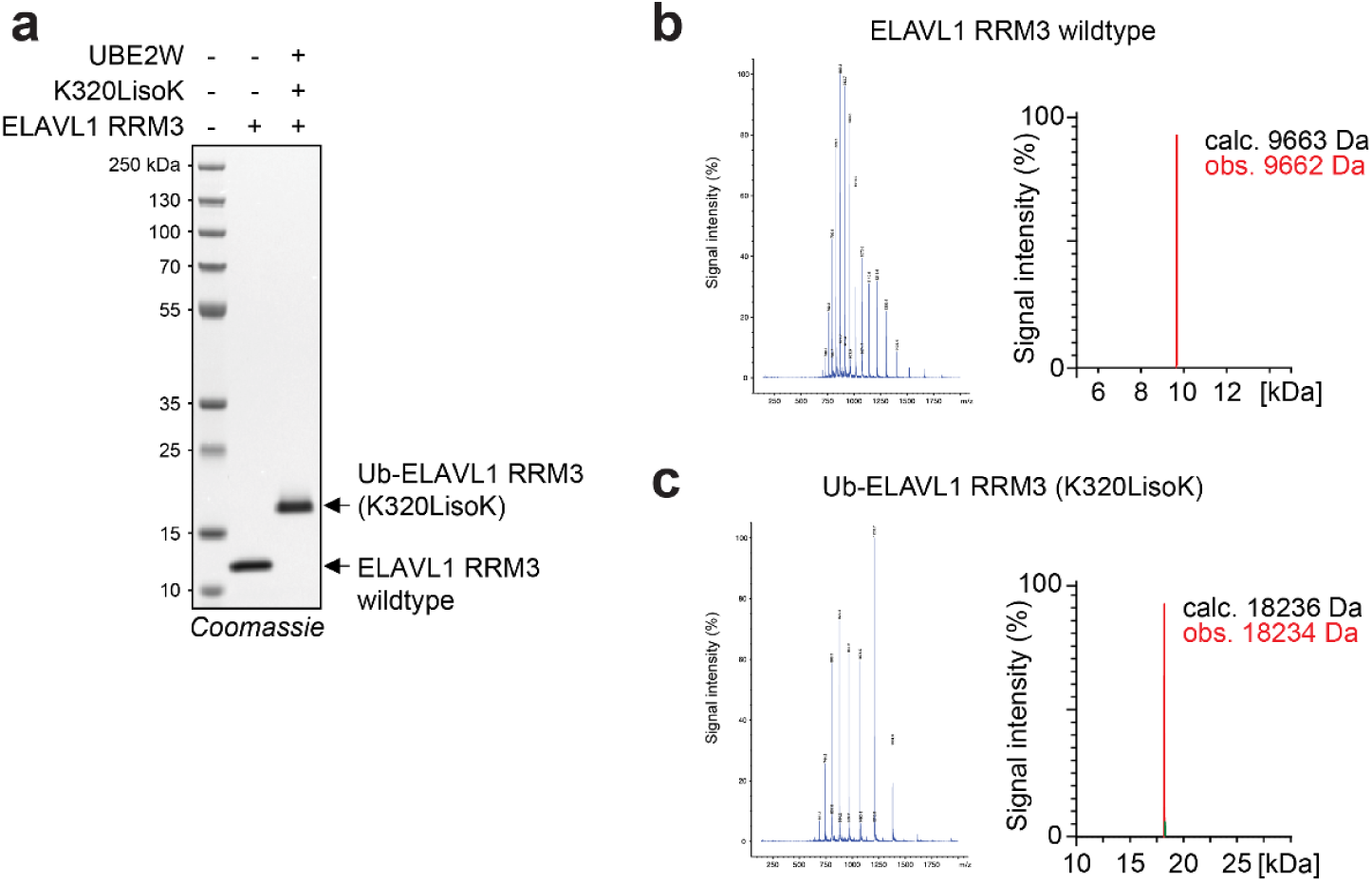
Confirmation of ELAVL1 RRM3 constructs A) Coomassie-stained SDS-PAGE of purified ELAVL1 constructs: Wildtype ELAVL1 RRM3 and ubiquitinated ELAVL1 RRM3 (K320LisoK). The cropped blot is shown in Fig. 5e. B-C) LC-MS measurement of purified B) ELAV1 RRM3 and C) Ub-ELAVL1 RRM3. Left: non-deconvoluted spectra, right: deconvoluted spectra.

## Supplementary tables and data

Table S1: Datasets in the human reference ubiquitinome

Table S2: The human reference ubiquitinome

Table S3: Pan-species ubiquitin proteomics datasets All_Experiments_nonHs: Meta-data from proteomics experiments in non-human species. Quantitative_Conditions: Meta-data of human quantitative proteomics data.

Table S4: Pan-species ubiquitin proteomics sites

Table S5: Ubiquitin site domain hotspots hotspotSummary: Summary of hotspots. hotspotSites: All ubi-sites found in hotspots. hotspotRegulatorySites: Hotspot ubi-sites with regulatory annotations in PhosphositePlus.

Table S6: Positional importance scores and features used for training

Table S7: Ubiquitin sites annotated to degradative and non-degradative functions in PhosphositePlus

Table S8: Ubiquitin sites in nuclear localisation signals or associated with protein activity

Table S9: Yeast chemical genetics experimental data conditionDescriptions: Descriptions of conditions used in growth screens. MutantDescriptions: Yeast strains used in growth screens. Sscores: S-scores and Q-values from chemical genetics experiments. primers: Primer sequences used for CRISPR-Cas9 generation of lysine mutants. donors: Donor sequences used for CRISPR-Cas9 generation of lysine mutants.

Table S10: Plasmids used in this study

Data S1: Ubi-site hotspots

Visualisation of all ubi-site hotspots with a hotspot found in a representative crystal structure.

