## Supplementary material for "Mapping the functional importance of site-specific ubiquitination across the human proteome": Data S1

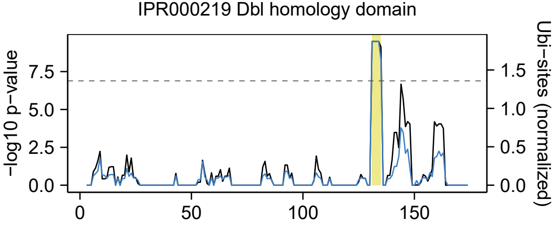

IPR000219 Dbl homology domain
−log10 p−value
Ubi−sites (normalized)
7.5
5.0
2.5
0.0
1.5
1.0
0.5
0.0
0
50
100
150


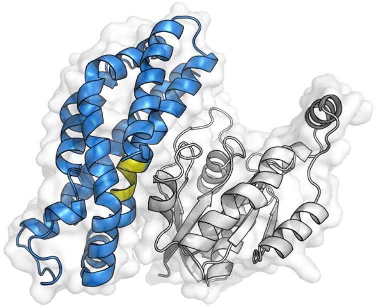

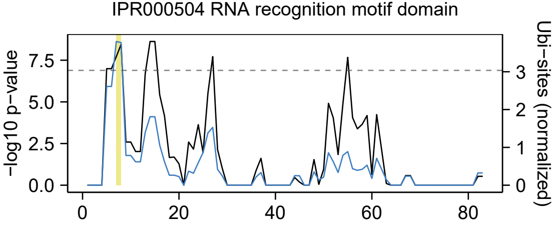

IPR000504 RNA recognition motif domain
-log10 p-value
Ubi-sites (normalized)
0.0
2.5
5.0
7.5
0
1
2
3
0
20
40
60
80


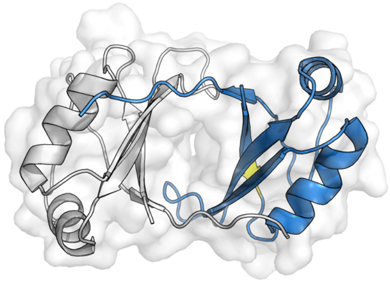

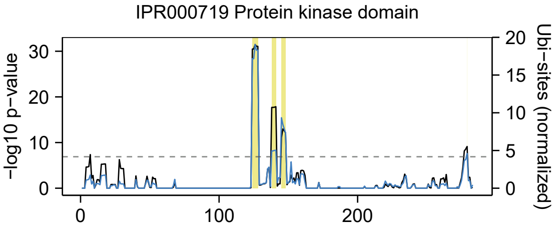
IPR000719 Protein kinase domain
-log10 p-value
Ubi-sites (normalized)
0
10
20
30
0
5
10
15
20
0
100
200

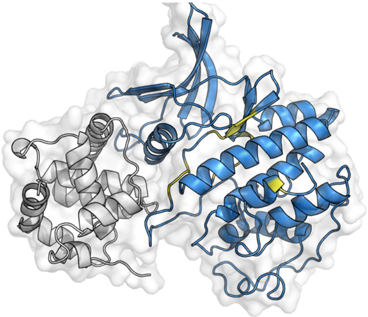

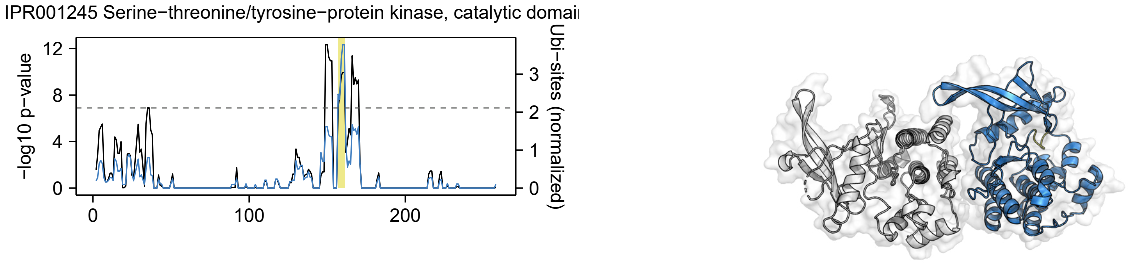
IPR001245 Serine−threonine/tyrosine−protein kinase, catalytic domai
-log10 p−value
Ubi−sites (normalized)
12
8
4
0
3
2
1
0
0
100
200

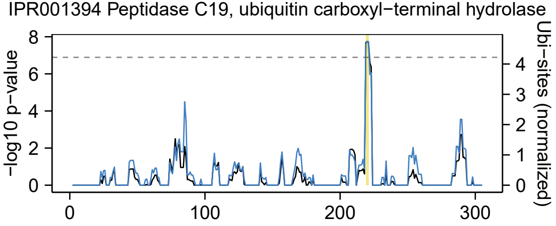
IPR001394 Peptidase C19, ubiquitin carboxyl−terminal hydrolase
-log10 p−value
Ubi−sites (normalized)
8
6
4
2
0
4
3
2
1
0
0
100
200
300

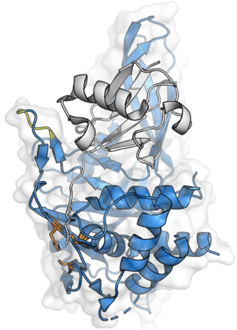

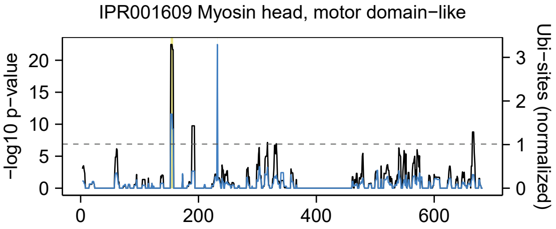
IPR001609 Myosin head, motor domain−like
-log10 p−value
Ubi−sites (normalized)
0
5
10
15
20
0
200
400
600
0
1
2
3

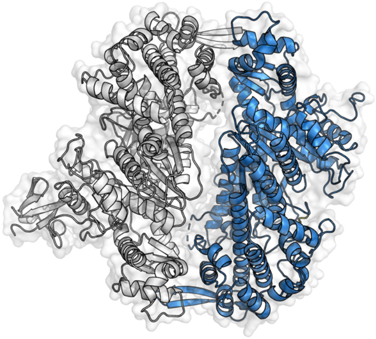

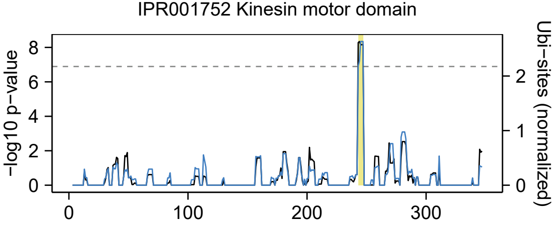

IPR001752 Kinesin motor domain
−log10 p−value
Ubi−sites (normalized)
0
2
4
6
8
0
1
2
0
100
200
300


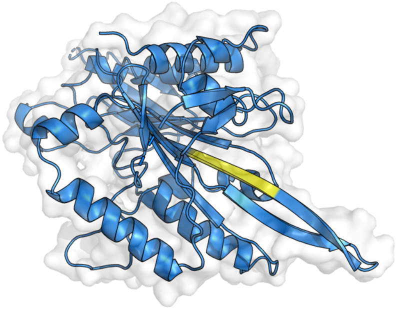

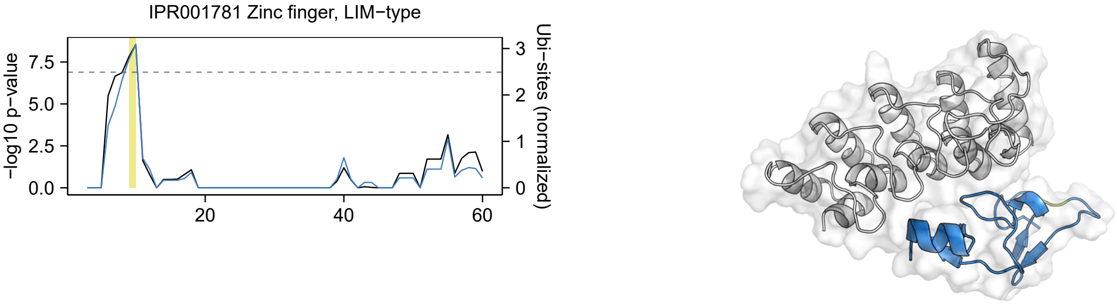
IPR001781 Zinc finger, LIM−type
−log10 p−value
Ubi−sites (normalized)
0.0
2.5
5.0
7.5
0
1
2
3
20
40
60

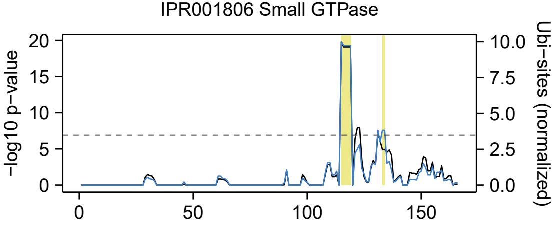

IPR001806 Small GTPase
-log10 p-value
Ubi-sites (normalized)
20
15
10
5
0
10.0
7.5
5.0
2.5
0.0
0
50
100
150


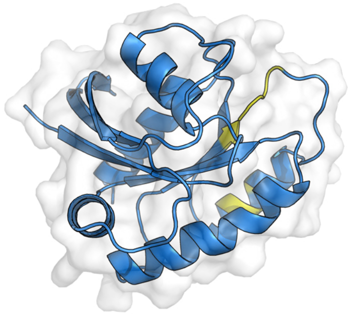

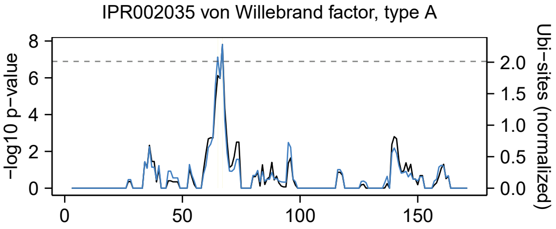
IPR002035 von Willebrand factor, type A
-log10 p-value
Ubi-sites (normalized)
8
6
4
2
0
2.0
1.5
1.0
0.5
0.0
0
50
100
150

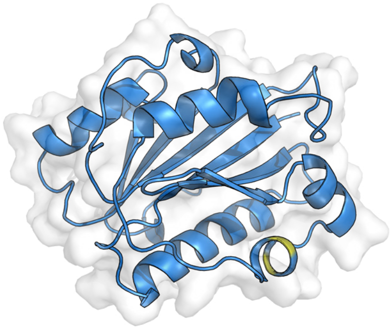

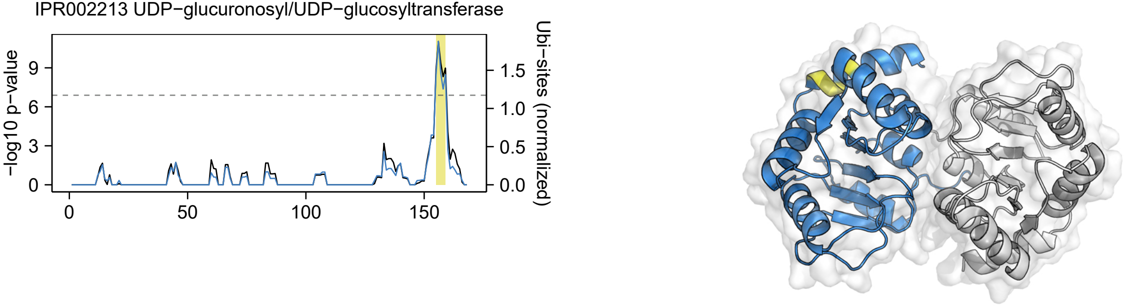
IPR002213 UDP−glucuronosyl/UDP−glucosyltransferase
-log10 p−value
Ubi−sites (normalized)
0
3
6
9
0.0
0.5
1.0
1.5
0
50
100
150

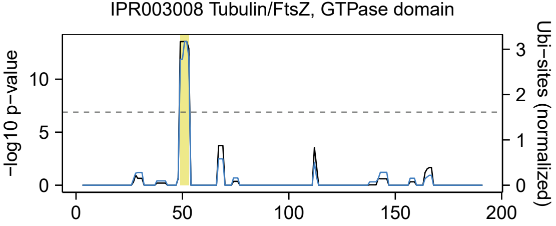
IPR003008 Tubulin/FtsZ, GTPase domain
-log10 p-value
Ubi-sites (normalized)
0
5
10
0
1
2
3
0
50
100
150
200

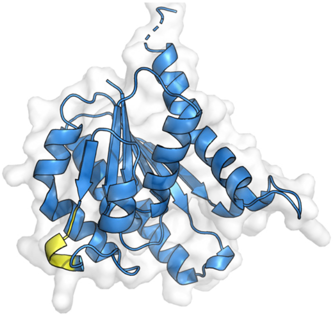

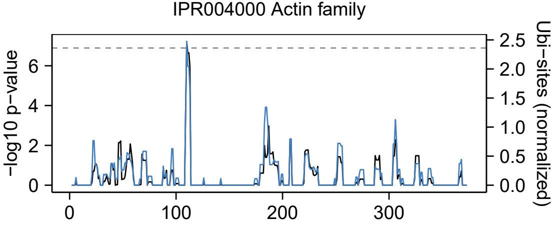
IPR004000 Actin family
-log10 p-value
Ubi-sites (normalized)
0
2
4
6
0.0
0.5
1.0
1.5
2.0
2.5
0
100
200
300

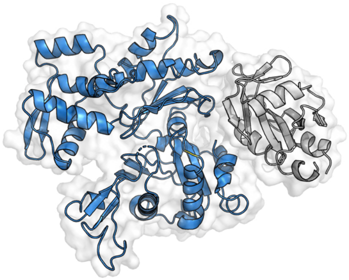

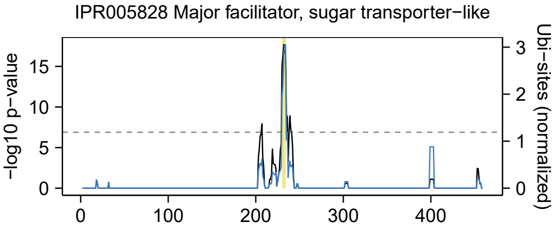
IPR005828 Major facilitator, sugar transporter−like
−log10 p−value
15
10
5
0
0
100
200
300
400
Ubi−sites (normalized)
3
2
1
0

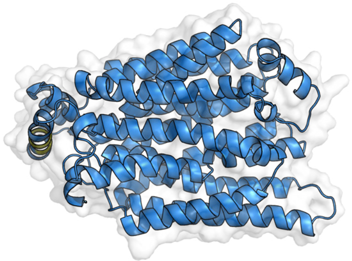

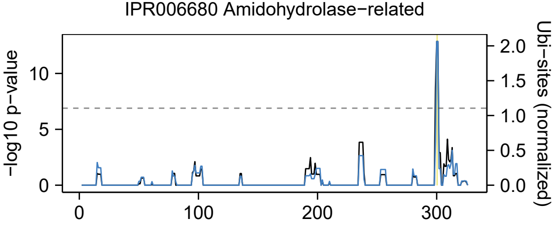

IPR006680 Amidohydrolase−related
−log10 p−value
Ubi−sites (normalized)
0
5
10
0.0
0.5
1.0
1.5
2.0
0
100
200
300


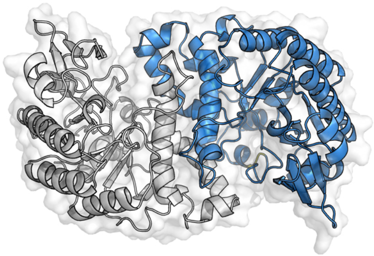

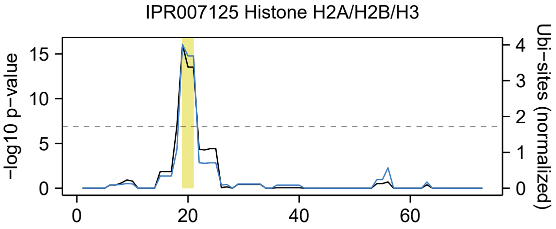

IPR007125 Histone H2A/H2B/H3
−log10 p−value
Ubi−sites (normalized)
15
10
5
0
4
3
2
1
0
0
20
40
60


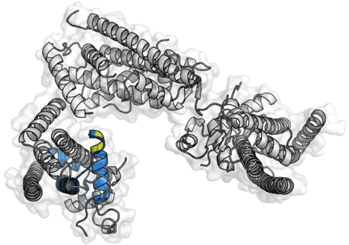

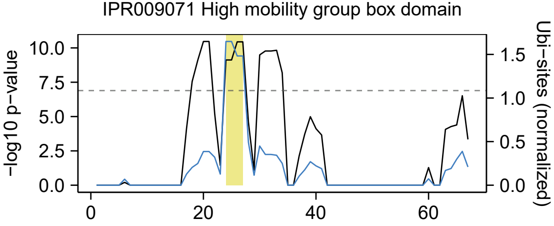
IPR009071 High mobility group box domain
-log10 p−value
Ubi−sites (normalized)
10.0
7.5
5.0
2.5
0.0
1.5
1.0
0.5
0.0
0
20
40
60
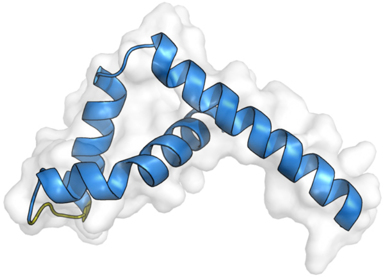

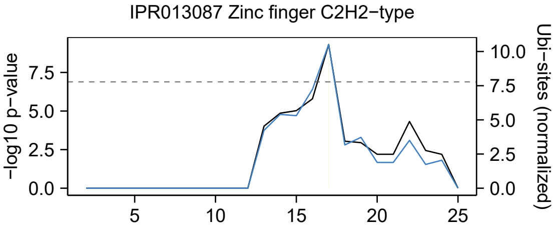
IPR013087 Zinc finger C2H2−type
−log10 p−value
Ubi−sites (normalized)
0.0
2.5
5.0
7.5
10.0
5
10
15
20
25

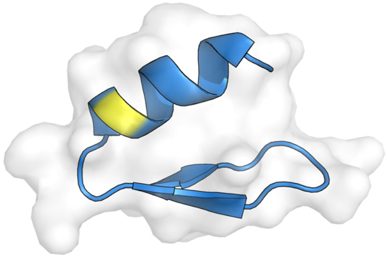

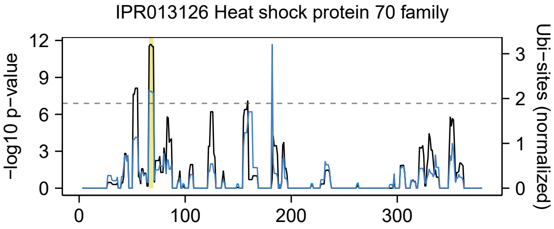

IPR013126 Heat shock protein 70 family
-log10 p−value
Ubi−sites (normalized)
12
9
6
3
0
3
2
1
0
0
100
200
300


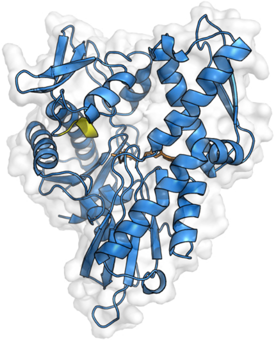

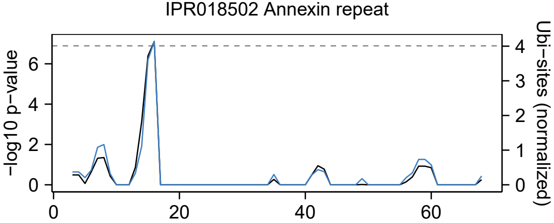
IPR018502 Annexin repeat
−log10 p−value
0
2
4
6
Ubi−sites (normalized)
0
1
2
3
4
0
20
40
60

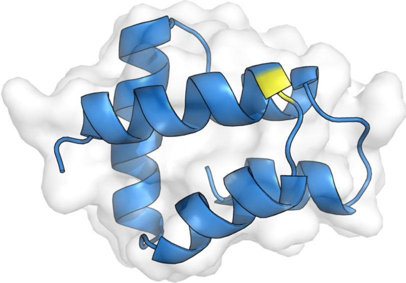

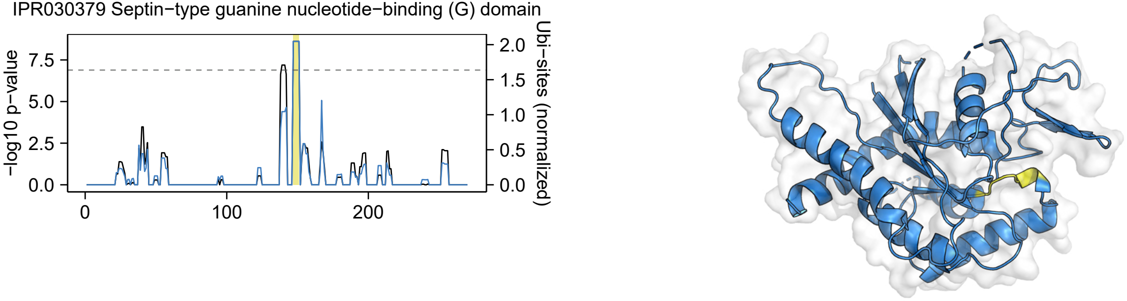
IPR030379 Septin−type guanine nucleotide−binding (G) domain
−log10 p−value
0.0
2.5
5.0
7.5
0
100
200
Ubi−sites (normalized)
0.0
0.5
1.0
1.5
2.0
